# NSD1 deposits histone H3 lysine 36 dimethylation to pattern non-CG DNA methylation in neurons

**DOI:** 10.1101/2023.02.17.528965

**Authors:** Nicole Hamagami, Dennis Y. Wu, Adam W. Clemens, Sabin A. Nettles, Harrison W. Gabel

## Abstract

During postnatal development the DNA methyltransferase DNMT3A deposits high levels of non-CG cytosine methylation in neurons. This unique methylation is critical for transcriptional regulation in the mature mammalian brain, and loss of this mark is implicated in DNMT3A-associated neurodevelopmental disorders (NDDs). The mechanisms determining genomic non-CG methylation profiles are not well defined however, and it is unknown if this pathway is disrupted in additional NDDs. Here we show that genome topology and gene expression converge to shape histone H3 lysine 36 dimethylation (H3K36me2) profiles, which in turn recruit DNMT3A and pattern neuronal non-CG methylation. We show that NSD1, the H3K36 methyltransferase mutated in the NDD, Sotos syndrome, is required for megabase-scale patterning of H3K36me2 and non-CG methylation in neurons. We find that brain-specific deletion of NSD1 causes alterations in DNA methylation that overlap with models of DNMT3A disorders and define convergent disruption in the expression of key neuronal genes in these models that may contribute to shared phenotypes in NSD1- and DNMT3A-associated NDD. Our findings indicate that H3K36me2 deposited by NSD1 is an important determinant of neuronal non-CG DNA methylation and implicates disruption of this methylation in Sotos syndrome.

**Highlights:**

- Topology-associated DNA methylation and gene expression independently contribute to neuronal gene body and enhancer non-CG DNA methylation patterns.
- Topology-associated H3K36me2 patterns and local enrichment of H3K4 methylation impact deposition of non-CG methylation by DNMT3A.
- Disruption of NSD1 *in vivo* leads to alterations in H3K36me2, DNA methylation, and gene expression that overlap with models of DNMT3A disorders.

## Introduction

Methylation of cytosines (mC) is an epigenetic mark that plays a critical role in transcriptional control in all mammalian cells. While all cells contain high levels of methylation at CG dinucleotides (mCG), neurons in the brain undergo a unique epigenomic reconfiguration in the early postnatal period (Lister et al., 2013) when the DNA methyltransferase DNMT3A deposits non-CG methylation across the genome. Non-CG methylation accumulates primarily at CA dinucleotides (mCA), reaching levels in many adult neuron types similar to that of mCG. These mCA sites are read out by the methyl-binding protein, MeCP2, to downregulate the activity of enhancers and tune the expression of genes with critical function in the nervous system (Boxer et al., 2020; Clemens et al., 2019; Gabel et al., 2015; Lagger et al., 2017).

Consistent with an essential role for the mCA pathway in the brain, disruption of either the deposition or readout of this mark has been implicated in neurodevelopmental disorders. Mutation of DNMT3A leads to the intellectual disability and overgrowth disorder, Tatton-Brown Rahman Syndrome (TBRS) (Sanders et al., 2015; Tatton-Brown et al., 2014), while loss or overexpression of MeCP2 leads to the severe postnatal onset neurological disorders Rett syndrome and MeCP2 duplication syndrome (MDS), respectively (Amir et al., 1999; del Gaudio et al., 2006; Van Esch et al., 2005). Epigenomic studies have demonstrated that neuronal mCA levels are globally reduced in mouse models of TBRS, leading to altered gene expression (Christian et al., 2020). These transcriptional changes overlap with the observed changes in gene expression that occur when readout of mCA is disrupted in models of Rett syndrome and MeCP2 duplication syndrome. Notably, exome sequencing studies of neurodevelopmental disorders have identified mutations in numerous genes that encode putative epigenetic regulators (Iossifov et al., 2014; Sanders et al., 2015; Satterstrom et al., 2020). In particular, mutation of the histone methyltransferase NSD1 leads to autism and other phenotypes in Sotos syndrome (Douglas et al., 2003; Kurotaki et al., 2002) that extensively overlap with those observed in TBRS (Tatton-Brown et al., 2017), but the impact of mutation of NSD1 on neuronal mCA has not been examined.

The level of mCA at gene regulatory elements determines the extent of repression by the mCA-MeCP2 pathway (Clemens et al., 2019), and thus it is critical to understand how mCA profiles across neuronal genomes are determined. Two independent phenomena have been associated with patterning mCA profiles, but how they interact to determine the ultimate levels of mCA at genomic regions has not been explored, and the molecular mechanisms that drive them are not known. In the first phenomenon, high levels of gene transcription during the early postnatal period (e.g. one to four weeks of age in mice) are associated with reduced binding of DNMT3A within the transcribed region of the gene and result in lower levels of mCA at this site in the mature brain (Stroud et al., 2017). The mCA levels determined by this transcription-associated effect can in turn impact whether a gene is subject to subsequent repression at the adult stage. At a mechanistic level, it is proposed that an alteration in chromatin structure or steric hinderance by the transcribing polymerase may mediate these effects, but no specific chromatin signal that is known to directly and negatively affect DNMT3A binding has been detected within genes. Therefore, the precise mechanism by which this “expression-associated” mCA protection occurs in neurons remains unclear.

Parallel studies have also discovered a second, large-scale phenomenon governing mCA levels (Christian et al., 2020; Clemens and Gabel, 2020; Clemens et al., 2019). Unlike mCG which is consistently high across chromosomes, mCA levels can vary substantially in broad, megabase-scale regions of the genome. This “regional mCA” is organized in part by topologically-associating domains (TADs) of chromatin folding. Genes and enhancers found within a TAD with a high mCA “set-point” will, on average, be more highly bound by DNMT3A in the early postnatal period and subsequently display higher mCA levels in the mature brain compared to genes and enhancers found within a TAD with a lower mCA set-point. Regional set-point mCA levels have important consequences for gene regulation in the adult brain, as enhancers found within high mCA TADs are reduced in activity leading to downregulated expression of target genes in that TAD. These genes are subsequently susceptible to dysregulation upon disruption of the mCA-MeCP2 pathway in disease (Clemens and Gabel, 2020; Clemens et al., 2019; Gabel et al., 2015). While these findings underscore the importance of regional mCA in gene regulation, it is not known what molecular mechanisms differentially recruit DNMT3A to TADs to determine the levels of mCA across these regions.

In this study we have investigated the mechanisms of DNA methylation patterning in the cerebral cortex of mice to build an integrated model of mCA deposition across genomic elements and uncover a key role for dimethylation of histone H3 lysine 36 by the Sotos syndrome protein NSD1 in this process. By analyzing gene expression and mCA levels across TADs and gene bodies, we show that regional mCA set-points and gene expression have independent, equal, and opposing contributions to the mCA profiles observed at genes and enhancers. By systematically profiling histone modifications previously not investigated in the early postnatal brain, we show that H3K4 methylation and H3K36 methylation are important predictors of DNA methylation at intergenic regions, genes, and regulatory regions. In particular, we find that the establishment of regional H3K36me2 levels, and selective depletion of this mark within genes, is a point of convergence that integrates both regional and gene expression-associated influences on mCA patterns. We show that loss of NSD1 disrupts H3K36me2 levels within TADs to alter DNMT3A targeting and mCA set-points, impacting mCA at genes and enhancers inside affected TADs. Notably, we find that alterations in DNA methylation and gene expression that result from disruption of NSD1 in the brain overlap epigenomic and transcriptomic changes in the mouse model of DNMT3A mutation in TBRS. Together, our findings present a unifying framework for mCA patterning in the neuronal genome and implicate disruption of neuron-specific DNA methylation as a shared site of molecular etiology across TBRS and Sotos syndrome.

## Results

### Regional set-point and gene expression additively and independently affect mCA at genes and enhancers

Both TAD-associated regional mCA set-points and gene expression-associated depletion of mCA have been shown to impact mCA levels across the neuronal genome (Clemens et al., 2019; Stroud et al., 2017), but whether they interact has not been assessed. For example, regional mCA levels could in fact be driven by expression of the genes within each TAD or gene expression could have different effects based on TAD context. Alternatively, they may independently influence mCA at genes (Figure 1A). Furthermore, the relative contribution of each to the ultimate levels of mCA has not been assessed. To explore how these two phenomena interact, we performed integrated analysis of the mouse cerebral cortex, examining TADs defined by HiC, gene expression measured by RNA-seq at two weeks of age, and adult DNA methylation assessed by whole-genome bisulfite sequencing at eight weeks of age. These time points are ideally suited for studying the epigenetic and transcriptional landscape at peak DNMT3A expression and mCA deposition in the early postnatal period, or after accumulation of adult mCA patterns, respectively. As previously reported (Clemens et al., 2019), genome-wide profiles of mCA in the adult exhibited differences in levels in broad, megabase-scale regions, with these fluctuations appearing far more strongly for mCA compared to mCG (Figure 1B, S1A). Further, robust depletion of mCA from expressed genes could be observed (Mo et al., 2015; Stroud et al., 2017). Visual inspection of genes in TADs with different mCA set-points (calculated as the levels of mCA in the TAD surrounding the gene, see *methods*) suggested that regional set-points influence mCA level at genes independently of gene expression (Figure 1B-E). Evidence of these effects can be observed at two genes (*Qk*, and *MeCP2*) that are found in TADs with different mCA set-points but have similar expression levels at two weeks (Figure 1C): *Qk*, which is found in a high mCA TAD, accumulates higher gene body mCA levels than the gene body of *MeCP2* which is found in a lower mCA TAD. Notably, the level of mCA depletion of the gene body for each gene relative to the regional mCA set-point is similar for the two genes, suggesting that gene expression determines the degree to which each gene is depleted compared to the regional methylation set-point. Quantification of mCA levels for TADs genome-wide further suggested that regional set-points for mCA are not tightly linked to the expression of genes within each TAD, as gene expression was robustly associated with the level of mCA depletion in each gene but showed little to no association with mCA in the remainder of the TAD surrounding the gene (Figure 1D). These effects were specific to mCA, as similar observations were not detectable for mCG (Figure S1B). To quantitatively assess the contributions of regional mCA and gene expression to gene mCA, we constructed a linear model using TAD-mCA levels and gene expression to predict mCA in genes. This analysis revealed that both mCA set-point levels and gene expression robustly correlate with the levels of mCA in genes, but that there is limited evidence of interaction between these two signals (Figure 1E). Notably, regional mCA and gene expression appeared to make approximately equal quantitative contributions to the overall level of mCA found within a gene, with each associated with up to about a two-fold difference in mCA across all genes, genome-wide (Figure 1D). Together these findings suggest that the regional mCA set-point predisposes a gene to high or low levels of mCA, and that the level of gene expression subsequently, and independently, influences the degree of mCA depletion below the TAD set-point.

**Figure 1.**
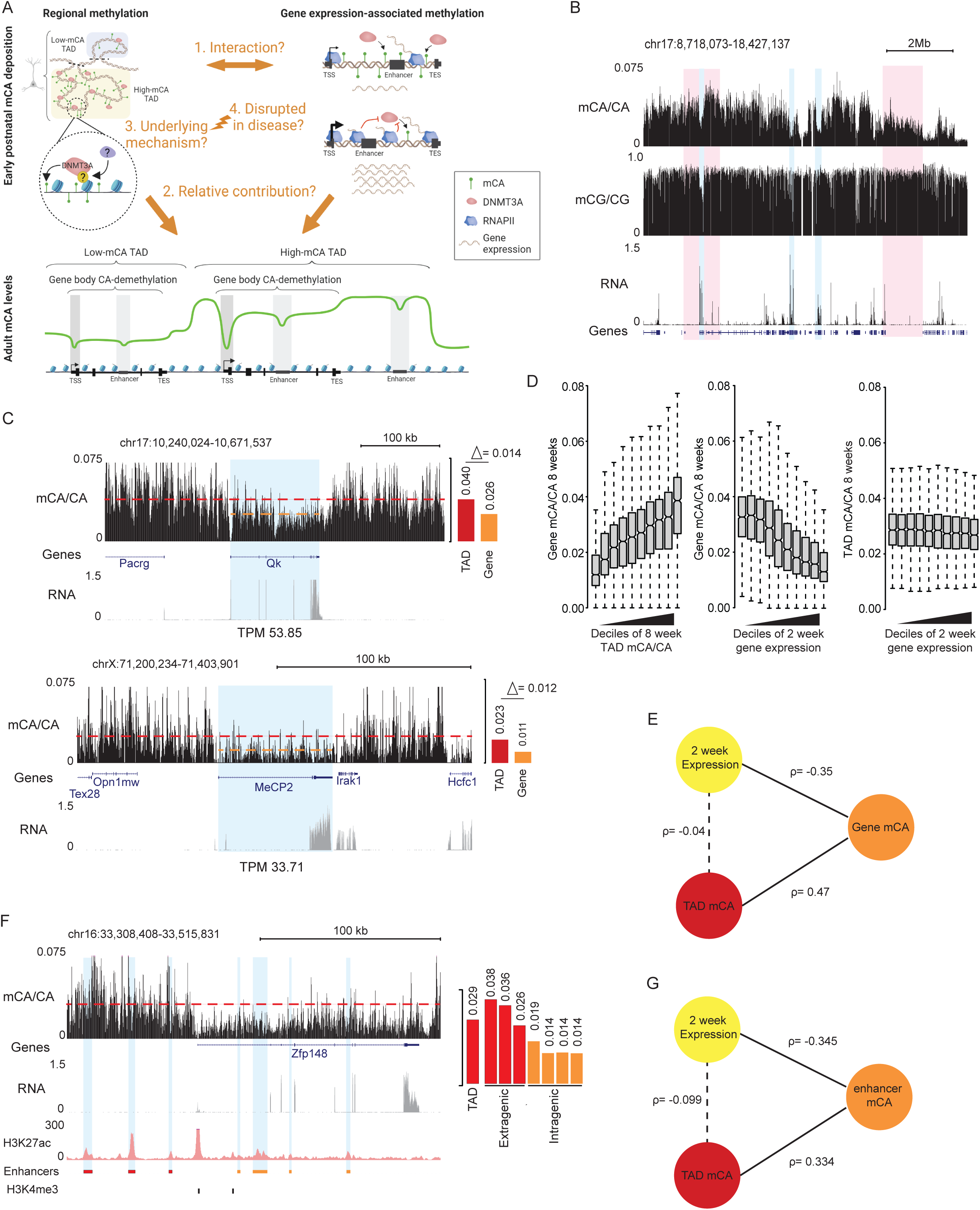
Regional mCA set-points and early postnatal gene expression converge to regulate neuronal mCA levels at genes and enhancers. (A) Left, model illustrating the deposition of non-CG methylation by DNMT3A to establish TAD-associated megabase-scale regions of high and low mCA. The underlying mechanism that guides DNMT3A to put down TAD-scale mCA, and whether this process is disrupted in disease is not known. Right, early postnatal gene expression antagonizes DNMT3A-mediated mCA deposition within gene bodies independently of TAD-directed mCA deposition. The degree to which these phenomena interact to determine mCA patterns, and how much each contributes to this process is not known (created with BioRender.com). (B) Genome browser view of total RNA-seq and DNA methylation from 2- and 8-week wildtype cerebral cortex, respectively. Genomic regions with high and low mCA levels are highlighted in red. Genes with varying levels of expression and depletion of mCA are highlighted in blue. In contrast to mCA, mCG shows little variation on a megabase-scale (see also, Figure S1). (C) Left, genome browser view of CA methylation (mCA) and RNA-seq from wildtype cerebral cortex at two genes, *Qk* and *MeCP2* (highlighted in blue), with similar expression levels but differing TAD mCA setpoints. Right, quantification of TAD mCA level (red) and gene body mCA level (orange). The average level of gene body mCA in the two genes differs while the amount of depletion of the gene body relative to the average level of mCA around the genes is similar. (D) Left, boxplots of adult mCA levels in the bodies of genes plotted for genes sorted into deciles of adult TAD mCA for each gene (TAD mCA is determined by the average level of mCA in the TAD in which the gene resides, see methods). Center, boxplot of adult mCA levels in the bodies of genes for genes sorted into deciles of gene expression at 2 weeks of age. Right, boxplot of adult TAD mCA levels for each gene sorted by deciles of gene expression at 2 weeks of age for each gene. Comparison of left and center plots indicates that regional TAD mCA set-point and 2 week gene expression in the early postnatal period are each associated with ∼2-fold variance in gene body mCA levels genome-wide. Right plot shows little to no association between 2 week expression and regional TAD mCA set-points. (E) Illustration of linear model and associated correlation coefficients observed between TAD mCA level, gene body mCA level, and gene expression. (F) Left, genome browser view of mCA, RNA-seq, and H3K27ac ChIP from wildtype cerebral cortex at *Zfp148* gene. Putative active enhancers (highlighted in blue) were defined as peaks that contain H3K27ac (red) and that do not overlap with promoter H3K4me3 peaks (black). Right, quantification of TAD mCA level (red), and intergenic (red) and intragenic (orange) enhancer mCA levels. Depletion of mCA at enhancers relative to the TAD mCA level is observed for intragenic enhancers within this highly expressed gene. (G) Illustration of linear model and associated correlation coefficients observed between TAD mCA level, intragenic enhancer mCA level, and gene expression. Data are from wildtype cerebral cortex at 2- and 8-weeks. 2-week RNA-seq data obtained from Lister et al., 2013. 8-week DNA methylation and H3K27ac ChIP-seq data obtained from Stroud et al., 2017 and Clemens et al., 2019.

Given that the levels of mCA at intragenic enhancers have been tightly linked to regulation of genes by MeCP2 (Clemens et al., 2019), we examined these regulatory sequences to determine how regional mCA set-points and early postnatal gene expression impact their ultimate mCA levels. Visual inspection (Figure 1F), quantitative analysis (Figure S1C), and linear modeling of all enhancers genome-wide (Figure 1G) indicated that intragenic enhancer mCA is robustly associated with both regional mCA set-points and gene expression, with depletion of mCA below the set-point being associated with higher gene expression. Thus, regional mCA and gene expression appear to additively and independently impact the ultimate levels of mCA at intragenic enhancers, which can in turn determine the degree of regulation at these sites by MeCP2 in the mature brain.

### Methylation of histone H3 lysine 36 predicts broad mCA patterns

We next investigated the molecular mechanism by which DNMT3A activity could be targeted to broad regions and then tuned at genes and enhancers to determine mCA levels at these sequences. Histone modifications have been shown to influence DNMT3A binding and activity on the genome (Fu et al., 2019; Li et al., 2021), but analysis of multiple histone modifications in the early postnatal brain has not detected any single modification that is a strong candidate for controlling mCA deposition by DNMT3A (Stroud et al., 2017). Notably, outside of the nervous system dimethylation of histone H3 lysine 36 (H3K36me2) has been implicated in recruiting DNMT3A to intergenic regions (Dukatz et al., 2019; Sankaran et al., 2016; Weinberg et al., 2019). Mutations in the PWWP domain of DNMT3A have been detected in TBRS patients suggesting that disruption of H3K36me2 binding by DNMT3A can drive neurodevelopmental deficits (Tatton-Brown et al., 2018). In addition, histone H3 lysine 4 methylation is thought to block activation of the enzyme at active gene-regulatory regions (Edwards et al., 2010; Ooi et al., 2007). However, these marks have not been studied for their impact on DNMT3A activity and deposition of mCA in the early postnatal brain. We therefore performed ChIP-seq analysis of H3K36 methylation (H3K36me2, and me3) and H3K4 methylation (H3K4me1, me2, and me3) in the cerebral cortex at two weeks of age. We assessed the degree to which these marks predict the binding of DNMT3A and patterns of mCA in the mature brain.

Examination of H3K36me2 across broad regions of the genome showed that this mark fluctuated on a megabase-scale, with profiles mirroring both DNMT3A binding across the genome at two weeks and subsequent patterns of mCA in the adult (Figure 2A). Cross-correlation analysis of H3K36me2 indicated that, like previously demonstrated for DNMT3A and mCA, this signal is more consistent within TAD structures than across TAD boundaries (Figure S2A). Aggregate analysis of H3K36me2 signal in and around genes and enhancers by quintiles of TAD mCA (Figure 2B), and quantitative genome-wide analysis (Figure 2C, S2B) revealed a robust correlation between this histone mark, DNMT3A binding, and mCA deposition at TADs, genes, and enhancers genome-wide, indicating that H3K36me2 patterns closely mimic the patterns expected for a signal that drives regional set-point mCA.

**Figure 2.**
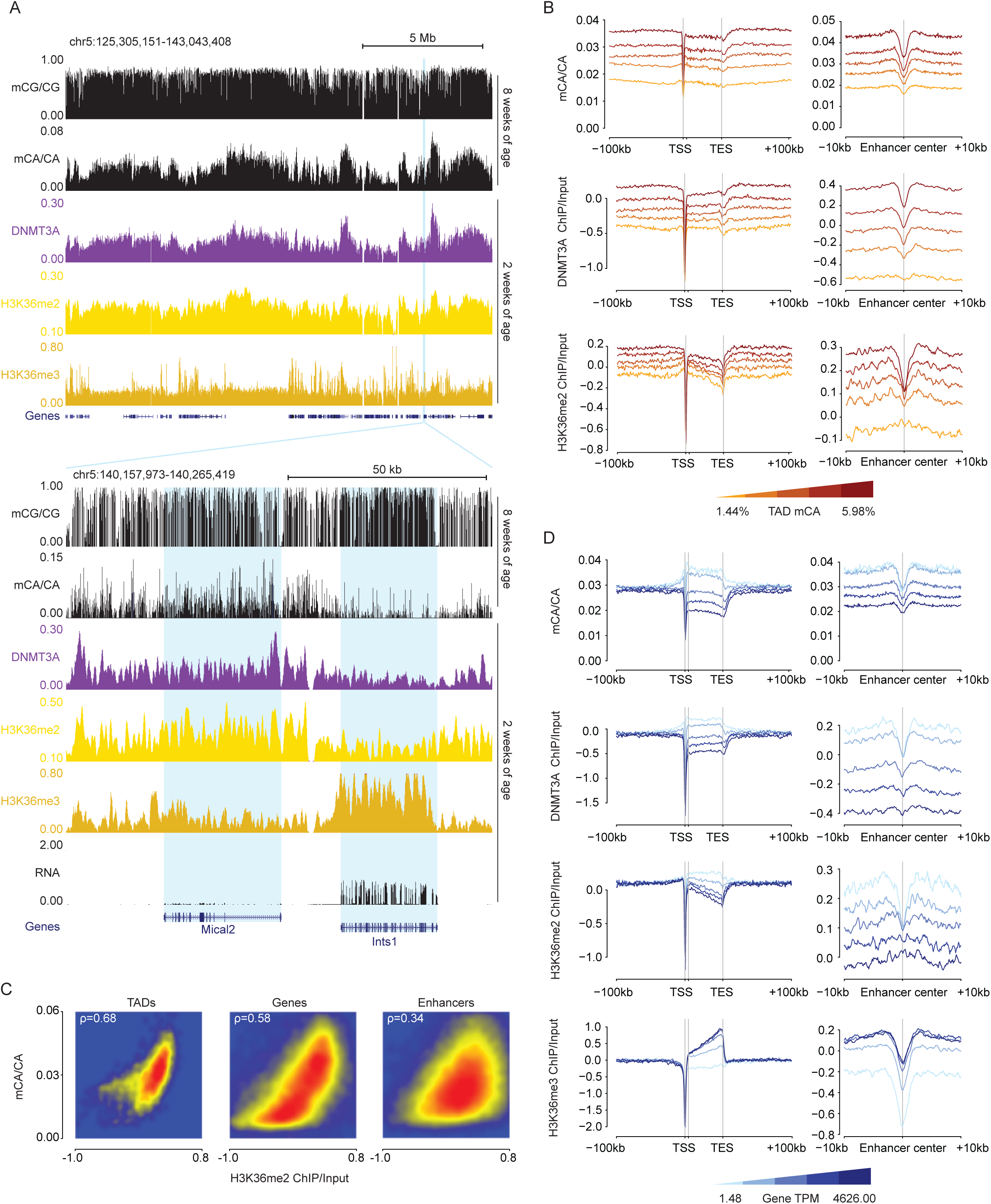
H3K36me2 profiles predict neuronal mCA in the postnatal cerebral cortex. (A) Top, broad genome browser view of DNA methylation, ChIP-seq, and total RNA-seq signal from wildtype cerebral cortex, illustrating megabase-scale fluctuations in H3K36me2 that correlate with DNMT3A binding and accumulation of mCA. Bottom, zoomed in genome browser view of two genes with different levels of expression, differential H3K36me2 and H3K36me3 ChIP signal, and corresponding differences in DNMT3A binding and mCA. (B) Aggregate mCA, DNMT3A, and H3K36me2 levels at genes and enhancers across quintiles of TADs sorted by regional mCA levels. (C) Smoothscatter plots of H3K36me2 ChIP and mCA/CA at TADs, enhancers, and genes, showing robust genome-wide correlations for these signals. (D) Aggregate plot of mCA, DNMT3A, H3K36me2, H3K36me3 levels at genes and enhancers across genes sorted by quintiles of RNA expression. Data are from wildtype cerebral cortex at 2- and 8-weeks. 2-week, n = 2-3 bioreplicates for H3K36me2 and DNMT3A ChIP-seq. 2-week RNA-seq data obtained from Lister et al., 2013. 2-week H3K36me3 ChIP-seq and 8-week DNA methylation data obtained from Stroud et al., 2017.

The conversion of H3K36me2 to H3K36me3 occurs in gene bodies and is strongly associated with the level of transcription through the gene (Wagner and Carpenter, 2012). Given the strong concordance between H3K36me2 and DNMT3A recruitment we considered that high expression of a gene in the early postnatal period may result in a relative increase of H3K36me3 that depletes H3K36me2 in genes. H3K36me2 has been reported to be a higher affinity binding site for DNMT3A than H3K36me3 (Dukatz et al., 2019; Weinberg et al., 2019; Xu et al., 2020), and such a conversion could explain the reduction of DNMT3A recruitment in highly expressed genes. Inspection of genes across the genome (Figure 2A) and aggregate analysis of genes as a function of expression levels (Figure 2D) supported this model, as highly expressed genes showed increased H3K36me3, depleted H3K36me2, and lower DNMT3A binding, with reduced subsequent mCA deposition at these sites in the adult brain. Notably, the expression associated depletion of H3K36me2 impacted mCA deposition at intragenic enhancers as well, with enhancers found within highly expressed genes showing similar depletion of this mark and subsequently lower mCA levels (Figure 2D). This finding supports a model in which H3K36me2 patterns across regions are established by broad deposition of H3K36me2, and depletion of this mark by conversion to the lower affinity DNMT3A binding site, H3K36me3, drives subsequent gene expression associated protection of gene bodies from methylation.

The association between H3K36me2 and DNA methylation in the brain was far stronger for mCA than for mCG, with mCG showing limited fluctuations on a megabase-scale across the genome (Figure 1B, 2A). In addition, we observed weaker correlations for mCG levels associated with levels of H3K36me2 in TADs, genes, and enhancers genome-wide (Figure S2C). Finally, mCG levels showed little dynamic range in genes as a function of their TAD mCA and gene expression levels (Figure S2D,E). Thus, in the context of the brain, H3K36me2 likely has its greatest impact on deposition of mCA by DNMT3A rather than on mCG.

### Focal depletion of mCA at regulatory elements is associated with histone H3 lysine 4 methylation

In addition to large-scale regional and gene body-associated mCA fluctuations, promoters of genes and distal enhancers often show focal depletion of mCA, suggesting that smaller-scale chromatin signals shape DNMT3A activity at these sites. H3K4 methylation is typically enriched at these regulatory sites when they are active, but it has not been systematically profiled in the early postnatal brain. *In vitro*, this histone mark can block the interaction of the DNMT3A ADD domain to the N-terminus of histone H3 resulting in inhibition of the enzyme at CG sites (Guo et al., 2015). Our ChIP-seq analysis of H3K4 mono-, di-and tri-methylation revealed a negative relationship between H3K4me2 and me3 and mCA deposition by DNMT3A, with high levels of these marks at promoters and enhancers that are depleted of DNMT3A binding and mCA levels (Figure 3A,B) and the strongest depletion occurring at promoters (Figure 3C, S3A). In contrast, H3K4me1 showed little to no association with mCA depletion compared to the higher H3K4 methylation states, suggesting that this mark does not strongly block ADD-mediated activation. This weaker effect of me1 on DNMT3A inhibition is supported by other studies showing that this mark does not strongly block DNMT3A ADD binding *in vitro* (Noh et al., 2015). mCG profiles showed similar relationships with H3K4 me2 and me3 as mCA (Figure S3B-D) indicating that these histone modifications likely play a similar, strong role in both mCA and mCG patterning. Notably, H3K4me1 was much more strongly associated with mCG depletion at regulatory elements compared to mCA, suggesting that there may be a differential effect of this histone mark on DNMT3A activity at these two dinucleotides. Together these findings support a model in which H3K4 di- and tri-methylation inhibit DNMT3A at active promoters and enhancers to block subsequent accumulation of neuronal mCA at these sites and raise potential differences in the importance of H3K4 mono-methylation in mCG and mCA patterning.

**Figure 3.**
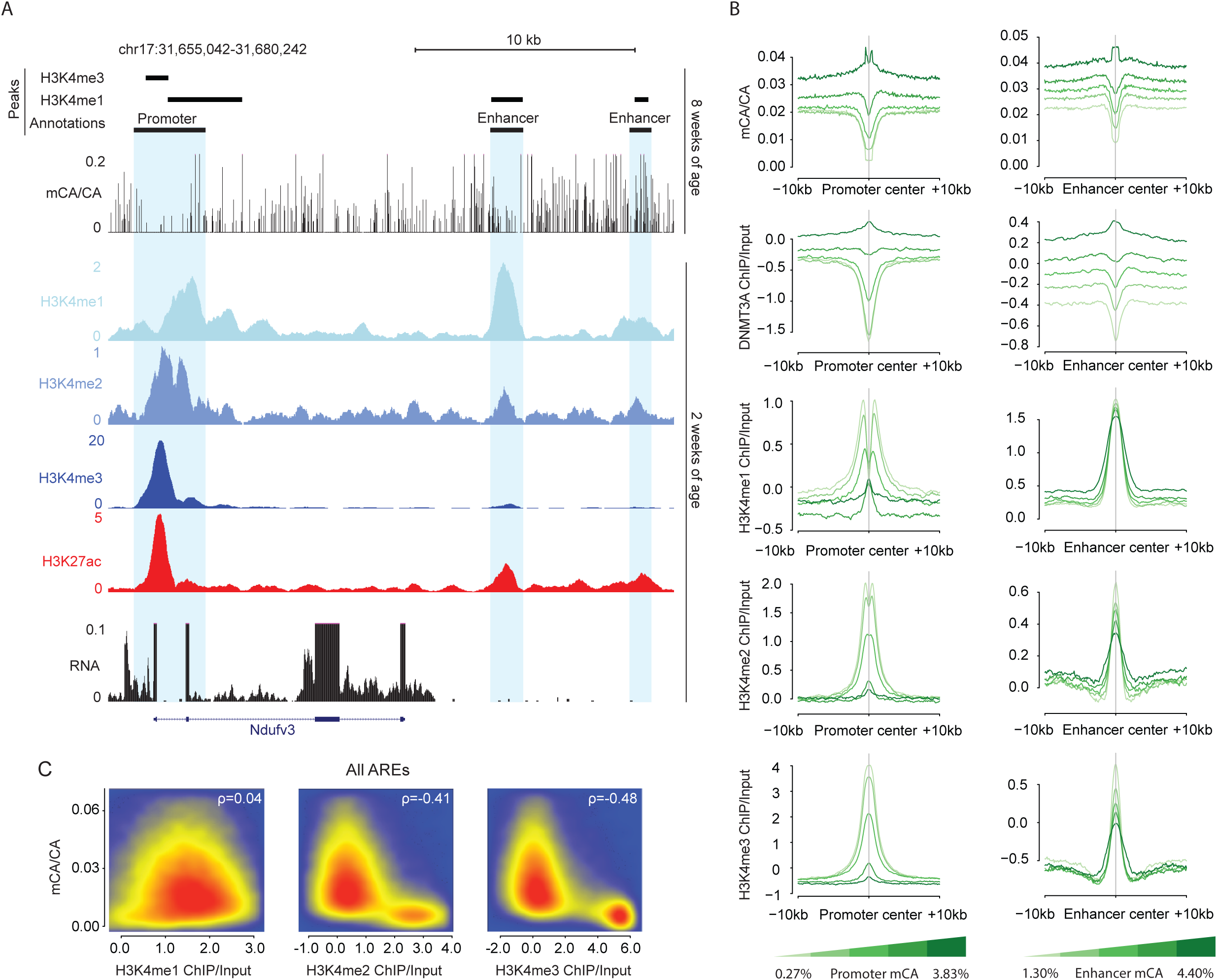
H3K4 methylation is negatively associated with DNMT3A-dependent neuronal mCA. (A) Genome browser view illustrating local enrichment of H3K4 methylation at active regulatory elements surrounding the *Nudfv3* gene and relative depletion of mCA at these sites. (B) Aggregate plots of mCA, DNMT3A, H3K4me1, H3K4me2, and H3K4me3 levels at transcription start sites (promoters) and enhancers sorted by quintiles of TSS or enhancer mCA levels. H3K4me2 and H3K4me3 show robust associations with mCA differences, with little differential signal for H3K4me1 detected. (C) Smoothscatter plots showing relative correlation of adult mCA levels with early postnatal H3K4 methylation marks at all active regulatory elements (AREs) across the genome (see *methods*). Data are from wildtype cerebral cortex at 2- and 8-weeks. 2-week, n = 2 for H3K4me1 and H3K4me2 ChIP-seq. 2-week RNA-seq data obtained from Lister et al., 2013. 2-week H3K4me3 ChIP-seq and 8-week DNA methylation data obtained from Stroud et al., 2017. H3K27ac ChIP-seq data obtained from Clemens et al., 2019.

### Integration of chromatin signatures accurately predicts mCA levels genome-wide

Given the robust associations of H3K36me2 and H3K4 methylation with DNMT3A-deposited mCA patterns, we next sought to compare the relative strength of these relationships to the signal for other chromatin states that have been previously profiled in the early postnatal brain (Stroud et al., 2017). We therefore carried out integrated analysis of these two marks with multiple ChIP-seq datasets measured in the cerebral cortex at two weeks of age. These data included histone modifications and RNA Polymerase II that display focal enrichment at active regulatory elements (H3K27ac; H3K4me1,2,3; RNAPII), histone marks and RNAP II modification states associated with gene bodies of highly expressed genes (H3K36me2,3; RNAPIIpS2), repressive histone modifications found broadly distributed across the genome (H3K27me2,3; H3K9me2,3), and histone variant binding profiles (H2Az; H3; H3.1,2). We assessed the correlation of each chromatin signal with gene expression, DNMT3A binding, and mCA deposition, examining this relationship at regulatory elements, genes, and intergenic/intragenic regions (Figure 4A). This analysis confirmed a strong negative correlation of H3K4 di- and tri-methylation with DNMT3A and mCA at regulatory elements that resembled similar relationships previously detected for Pol II binding at promoters (Stroud et al., 2017). Intragenic signals associated with high gene expression (Pol II pS2, H3K36me3) showed negative correlations with DNMT3A and mCA. Notably, at all sites examined, we found that H3K36me2 consistently showed a positive correlation with DNMT3A and mCA deposition, while showing a negative correlation with gene expression. To examine the degree to which the chromatin signatures profiled to date can explain mCA deposition patterns, we trained a random forest algorithm to predict mCA levels with these data using 5-fold cross-validation (Dietterich, 1998; Liaw and Wiener, 2002; Stone, 1974). The algorithm accurately predicted mCA for both intergenic and intragenic regions genome-wide (Figure 4B). Notably this random forest approach outperformed a simple linear model (Figure S4A), suggesting that DNMT3A binding could not be predicted solely by any one chromatin factor or any simple combination of factors and implying a more complex, nonlinear relationship between these factors and mCA deposition. To assess the contribution of each ChIP signal to the prediction accuracy of the algorithm we performed feature importance and feature elimination analyses (Figure 4C,D, S4C). Notably, this revealed H3K36me2 as one of the strongest contributors to accurate prediction of mCA levels across regions genome-wide. Analysis of regulatory regions alone revealed a larger signal for H3K4me3 at promoters (Figure S4B), further implicating this mark as a determinant of mCA at focal regulatory sites of active genes. Together these findings further support key roles for H3K4me3 and H3K36me2 in patterning neuronal methylation, with H3K36me2 showing a particularly strong signal across the genome regardless of location.

**Figure 4.**
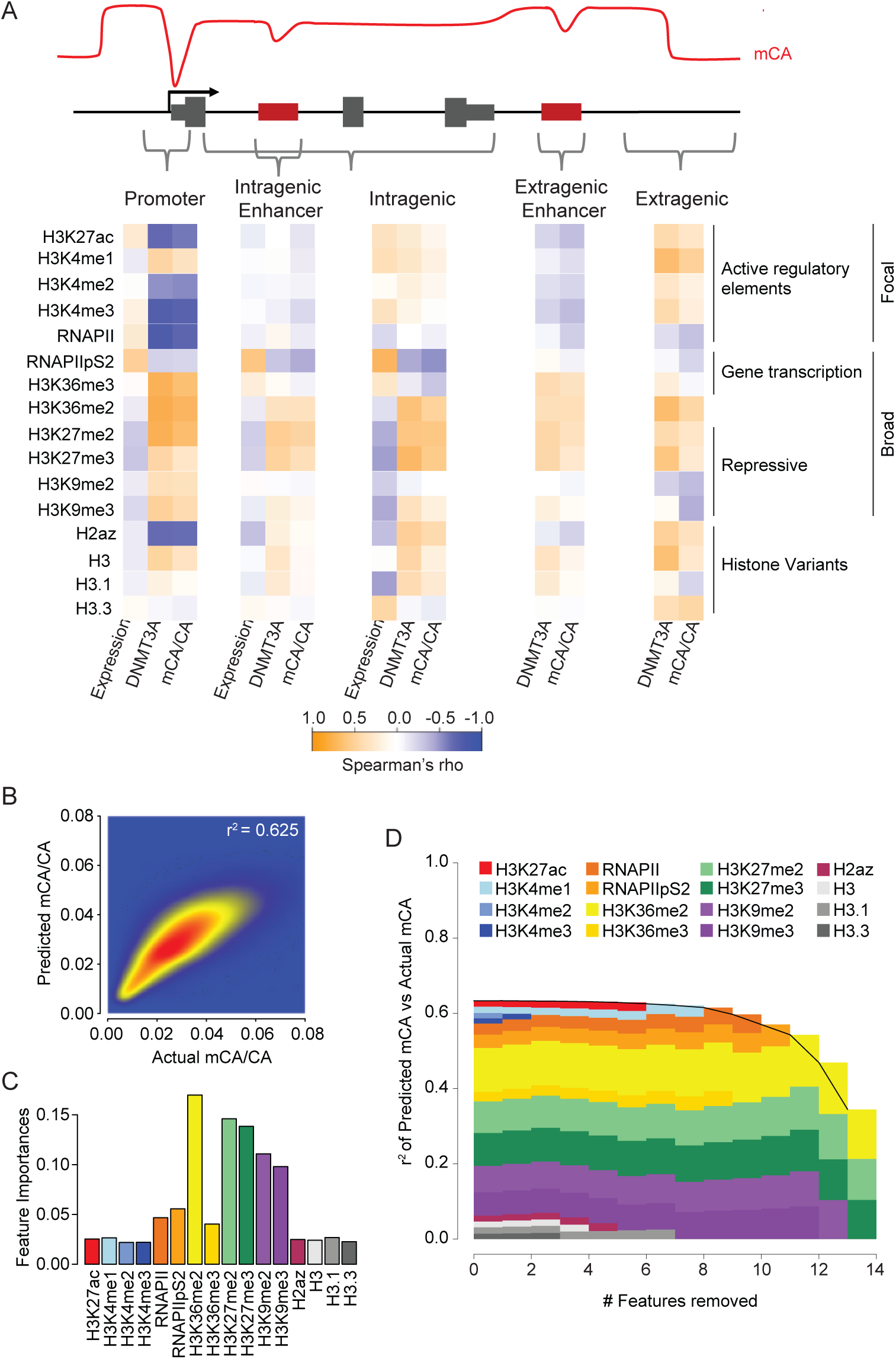
Comparative analysis of the association between chromatin states, DNMT3A binding and mCA deposition across the neuronal genome. (A) Correlation of histone and RNA polymerase II modification states with DNMT3A binding, mCA, and gene expression across different kilobase-scale genomic regions. (B) Random forest classifier predicted mCA versus actual mCA levels genome-wide. (C) Feature importance analysis indicating the relative contribution of each chromatin signature to the predictive accuracy of mCA levels by the random forest classifier. (D) Feature elimination analysis reveals key marks required for accurate prediction of mCA levels across the genome by the random forest classifier. Data are from wildtype cerebral cortex at 2- and 8-weeks. 2-week, n = 2-3 bioreplicates for H3K36me2, H3K4me1, H3K4me2, and DNMT3A ChIP-seq. 2-week RNA-seq data obtained from Lister et al., 2013. Remaining 2-week chromatin mark ChIP-seq and 8-week DNA methylation data obtained from Stroud et al., 2017, (see *methods*).

### Disruption of NSD1 in neurons leads to reductions of H3K36me2 and loss of mCA in broad domains

To directly test the importance of H3K36me2 in DNMT3A targeting and mCA patterning, we sought to disrupt H3K36me2 deposition and assess its impact on neuronal methylation. Previously, we have demonstrated that deposition of mCA by DNMT3A can be observed in primary cortical neurons isolated at embryonic day 14.5 (E14.5) and cultured *in vitro* for 12 or more days (Christian et al., 2020). Having used this system to examine the effects of NDD-associated DNMT3A mutations on mCA deposition, we sought to employ it to manipulate H3K36me2 profiles and evaluate the effects on DMMT3A-mediated mCA patterning. Given the shared cognitive and behavioral deficits upon NSD1 mutation in Sotos syndrome and DNMT3A mutation in TBRS we selected NSD1 for disruption in our cultures by lentiviral-mediated RNAi knockdown. Two shRNAs were selected and validated for NSD1 knockdown (Figure S5A) and compared to control in ChIP-seq and WGBS analysis. Viral transduction was performed at Day *in vitro* (DIV) 1 followed by ChIP-seq of H3K36me2, DNMT3A binding at DIV 12, and assessment of subsequent mCA and mCG levels by WGBS analysis of DNA isolated at DIV 18. Inspection of ChIP-Seq and DNA methylation profiles in control neurons further supported a strong association between H3K36me2, DNMT3A binding, and mCA deposition (Figure S5B-D), with the profiles of these signals closely mirroring each other across the genome. To examine the effects of NSD1 knockdown on H3K36me2 and DNMT3A targeting genome-wide we analyzed each ChIP-seq signal using differential count analysis by edgeR (Nikolayeva and Robinson, 2014), identifying significantly changed 10kb windows across the genome. We detect 20,203 10kb windows with significantly altered H3K36me2 levels upon knockdown of NSD1. Analysis of DNMT3A ChIP-seq signal and WGBS in these regions showed changes in DNMT3A binding and mCA that are highly concordant with observed alterations in H3K36me2 (Figure 5A,B). To extend our analysis beyond the subset of regions called as significant and assess the effect of NSD1 knockdown genome-wide, we examined changes in all 10kb windows across the genome. This analysis revealed a strong global concordance between changes in H3K36me2 signal, DNMT3A signal, and mCA (Figure 5C). These effects were seen in aggregate analysis, as well as when the two NSD1 shRNAs were independently interrogated (Figure S5E), demonstrating the role of H3K36me2 in targeting DNMT3A to the neuronal genome.

**Figure 5.**
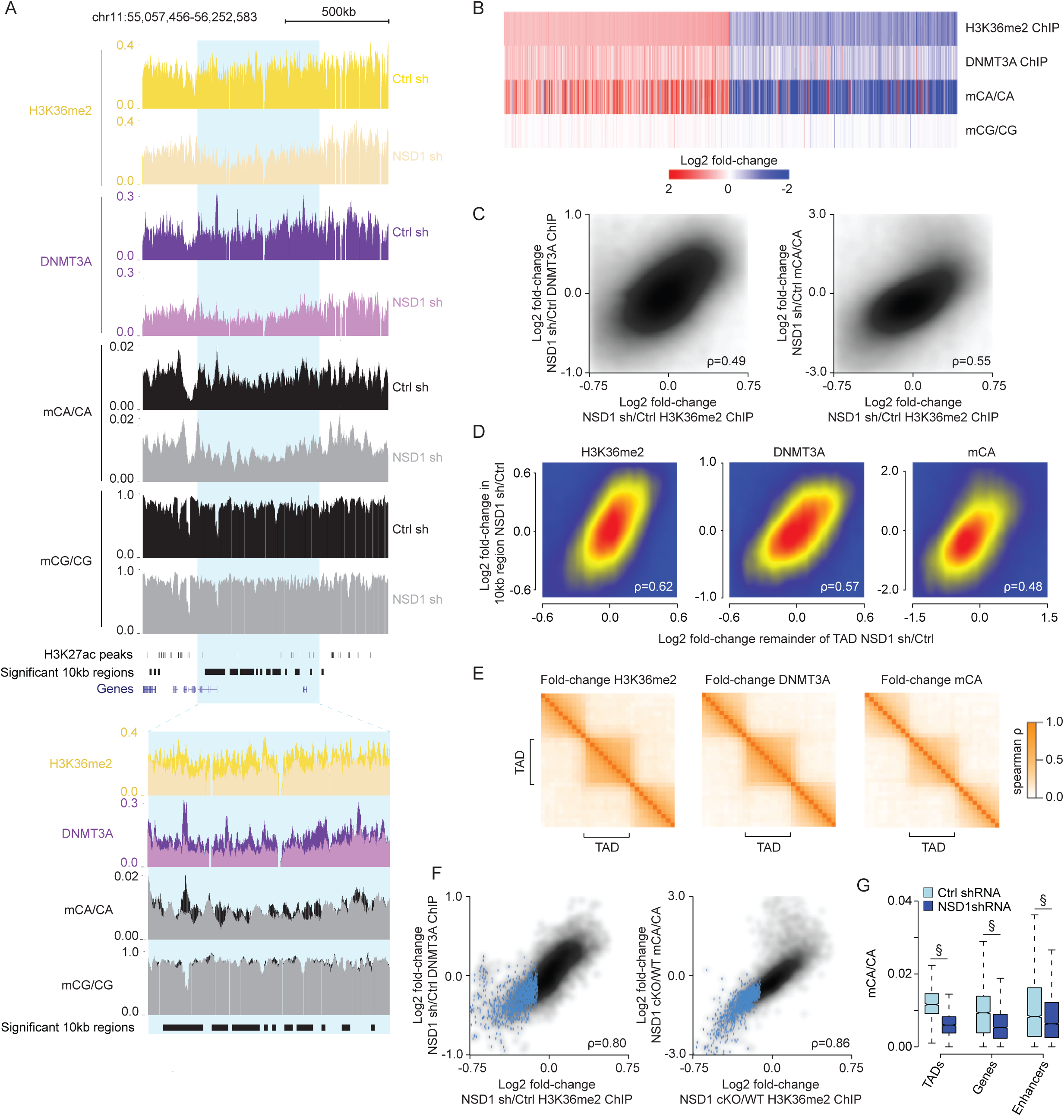
NSD1-mediated H3K36me2 is required for TAD-scale DNMT3A targeting and mCA deposition in postmitotic neurons. (A) Genome browser view of DIV 12 ChIP-seq and DIV 18 DNA methylation from primary cortical neurons (PCN). A representative TAD with significantly reduced H3K36me2 upon NSD1 knockdown is highlighted in blue. Overlap of multiple 10kb bins detected as significantly reduced for H3K36me2 with this significantly altered TAD illustrates strong concordance of changed signals within TADs. (B) Fold-changes of H3K36me2, DNMT3A, and DNA methylation in shNSD1 transduced PCNs at all 10kb regions identified as having significantly altered H3K36me2 signal by edgeR (FDR<0.1). (C) Smoothscatter analysis of fold-change of H3K36me2 and fold-change of DNMT3A (left) or mCA (right) for each 10kb region in the genome. (D) Smoothscatter analysis of changes in H3K36me2, DNMT3A, or mCA for 10kb regions of the genome compared to the TAD in which each region resides. (E) Cross-correlation analysis of fold-changes in H3K36me2, DNMT3A, and mCA upon knockdown of NSD1 for regions inside and outside of TADs across the genome. The fold-change for each signal was assessed for 10 genomic windows within each TAD as well as for 10 equally sized windows on either side of each TAD. The correlation between the fold-changes in these windows was assessed for all TADs genome-wide and plotted based on position relative to the TAD (see *methods*). The higher correlation of regions found within the same TAD compared to regions across TAD boundaries indicates that regions found within the same TAD are concordantly affected upon loss of NSD1. (F) Smoothscatter comparing changes in H3K36me2 to changes in DNMT3A (left) or mCA (right) for each TAD in the genome. TADs with significantly reduced H3K36me2 by edgeR (FDR<0.1) are highlighted in blue. (G) Boxplots of mCA levels in shCtrl and shNSD1 PCNs at TADs with significantly reduced H3K36me2 (edgeR, FDR <0.1) and kilobase-scale genomic elements that reside within these TADs. (§p < 10^-15^, Wilcoxon test). Data are from PCNs infected with shCtrl or shNSD1 after 1 Day *in vitro* (DIV) and collected at DIV 12 and DIV 18 for ChIP-seq and WGBS, respectively. Per time point: n = 2-4 bioreplicates for H3K36me2, DNMT3A ChIP-seq, and DNA methylation. TADs were derived from Hi-C analysis of cortical neurons (Bonev et al., 2017).

Given that we have detected that H3K36me2, DNMT3A, and mCA are organized by TADs, we wanted to assess if NSD1-mediated H3K36me2 deposition is targeted to TADs and determine if disruption of this enzyme therefore impacts mCA deposition across entire TADs. To investigate this, we first analyzed changes in 10kb windows, examining if they are correlated with changes in broader TAD regions in which they are found. Indeed, we observe that the changes in each 10kb window is associated with the remainder of the TAD in which the 10kb window resides (Figure 5D), suggesting that changes in 10kb windows reflect a larger scale effect on the TAD. Furthermore, cross-correlation analysis shows that alterations in H3K36me2, DNMT3A, and mCA upon NSD1 knockdown were organized by TADs (Figure 5E). Consistent with TADs being the units targeted by NSD1 for H3K36me2 and mCA deposition, genome-wide analysis of changes in the levels of H3K36me2, DNMT3A, and mCA showed extremely strong concordance (Figure 5F). These findings indicate that the intrinsic scale of H3K36me2 targeting by NSD1 is at the level of TADs.

Our analysis above (Figure 2) suggests that the level of H3K36me2 in TADs can influence the level of this mark and subsequent mCA deposition in smaller scale genes and regulatory elements that reside in each TAD. We therefore assessed how mCA is affected at enhancers and genes in TADs with reduced H3K36me2 signal upon knockdown of NSD1. We observe reduced mCA at these sites when they are found within TADs that display significantly reduced H3K36me2 signal, indicating that NSD1-mediated deposition of H3K36me2 in TADs helps to determine the level of mCA at these important sites of genes regulation (Figure 5G). Together these analyses indicate that NSD1 activity is required to maintain H3K36me2 patterns in TADs in postmitotic neurons, and that this H3K36me2 profile directs regional mCA set-points to impact the levels of mCA at regulatory elements and genes across the genome.

### Shared epigenomic and transcriptomic effects of NSD1 and DNMT3A disruption *in vivo*

Having detected disruption of H3K36me2 and mCA upon acute depletion of NSD1 in cultured neurons, we next sought to assess how loss of NSD1-mediated H3K36me2 deposition in the brain affects the neuronal methylome and manifests as alterations in chromatin and gene expression. In particular, our *in vitro* findings that loss of NSD1 leads to altered DNMT3A binding and mCA deposition suggest that there might be direct mechanistic overlap between the effects of NSD1 disruption and those observed upon DNMT3A mutation. Such shared changes would be candidates to explain the phenotypic overlap observed between Sotos syndrome and TBRS.

To carry out *in vivo* studies, we generated and analyzed NSD1 conditional knockout (cKO) mice in which deletion of exon 3 in the brain (Oishi et al., 2020) was driven by the expression of *Nestin*-cre in neural progenitors (Figure S6A). Validation of the NSD1 cKO mouse showed efficient removal of the targeted exon selectively in the brain and resulted in disruption of the NSD1 mRNA via altered splicing and early truncation of the coding sequence, indicating that the majority of the NSD1 gene product was disrupted in the brain (Figure S6B-D).

To determine if disruption of NSD1 *in vivo* alters the neuronal DNA methylation pathway, we first assessed changes in H3K36me2 and DNMT3A at two weeks of age, during peak DNMT3A expression, to determine the impact on targeting of the enzyme. We performed ChIP-seq on the cerebral cortex of NSD1 cKO and control mice for this analysis, utilizing the ChIP-Rx procedure (Orlando et al., 2014) to allow us to assess if global levels of these signals are altered in the mutant. Differential count analysis by edgeR revealed similar TAD-level changes in H3K36me2 and DNMT3A that we observed in culture (Figure 6A,B), with reductions in TAD H3K36me2 strongly correlating with loss of DNMT3A binding. We then performed base-pair resolution whole genome bisulfite sequencing in eight week adult cortex to assess the impact of NSD1 loss on DNA methylation in the mature brain. Analysis of mCA detected changes concordant with altered DNMT3A targeting, including lower mCA levels in TADs that showed reduced H3K36me2 at two weeks of age (Figure 6B). A concomitant reduction in mCA at genes and enhancers found within affected TADs was observed (Figure 6C), consistent with the impact of reducing the mCA set-point of these broad regions. In addition to a significant reduction of mCA at an array of TADs across the neuronal genome, we noted a trend toward lower mCA levels globally, with total mCA levels showing a reduction by approximately 10%. Analysis of ChIP-seq yields by ChIP-Rx normalization indicated a similar trend toward global loss of H3K36me2 (7%) upon loss of NSD1 (Figure S6E). Compared to mCA, mCG levels were largely maintained globally and across broad genomic elements in which H3K36me2 was lost, with only subtle effects observed that mirrored changes in mCA (Figure 6A, S6F). Together these results indicate that mCA levels in a subset of TADs in the genome are particularly sensitive to loss of NSD1 in the brain, and that NSD1 contributes to global H3K36me2 and subsequent DNMT3A-mediated mCA levels genome-wide.

**Figure 6.**
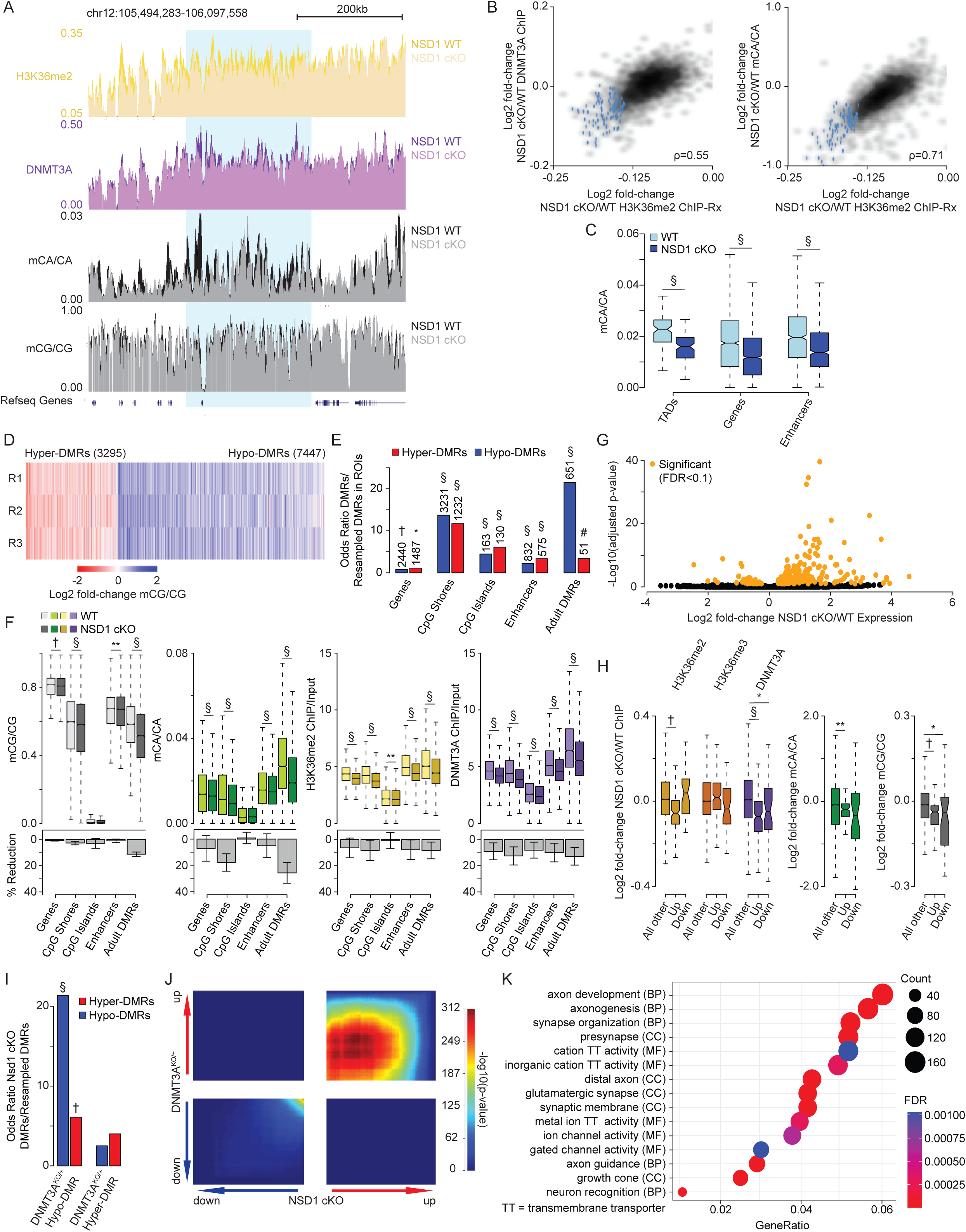
Brain-specific loss of NSD1 *in vivo* leads to epigenetic and transcriptional dysregulation that overlap DNMT3A mutants. (A) Genome browser view of two week ChIP-Rx and eight week DNA methylation from NSD1 cKO and wildtype cerebral cortex. A representative TAD with significantly reduced H3K36me2 is highlighted in blue. (B) Smoothscatter showing changes in H3K36me2 and changes in DNMT3A (left) or mCA (right) for each TAD. TADs with significantly reduced H3K36me2 by edgeR (FDR<0.1) are highlighted in blue. (C) Boxplots showing mCA levels at TADs with significantly reduced H3K36me2 (edgeR, FDR <0.1) and kilobase-scale genomic elements that reside within these TADs in NSD1 cKO and wildtype cerebral cortex. (Wilcoxon test). (D) Fold-change of mCG at CG-DMRs called in the NSD1 cKO cortex across three bioreplicates. (E) Odds ratio of overlap of NSD1 cKO CG-DMRs with kilobases-scale genomic regions, with the observed number of overlapping regions denoted above(Fisher’s exact test, observed versus background estimated from resampled DMRs, see *methods*). (F) Top, DNA methylation, H3K36me2 levels, and DNMT3A binding in NSD1 cKO versus wildtype cortex across kilobase-scale genomic regions. Bottom, mean and SEM percent reduction across genomic regions. (Wilcoxon test). (G) Volcano plot of fold-changes in gene expression of NSD1 cKO versus wildtype cortex total RNA-seq. Genes called as significant (FDR <0.1) by DESeq2 are highlighted in orange. (H) Fold-changes of H3K36me2, H3K36me3, DNMT3A, and DNA methylation at genes identified as significantly dysregulated (DESeq2, FDR <0.1) by RNA-seq in the NSD1 cKO. (Wilcoxon test). (I) Odds ratio of overlap of NSD1 cKO CG-DMRs with DNMT3A^KO/+^ CG-DMRs (Christian et al., 2020). (Fisher’s exact test, observed versus background estimated from resampled DMRs, see *methods*). (J) Rank-rank hypergeometric overlap (RRHO) (Cahill et al., 2018; Plaisier et al., 2010) of transcriptome-wide gene expression changes in the cerebral cortex of NSD1 cKO versus DNMT3A^KO/+^ mice (Christian et al., 2020). (K) Top five Gene Ontology terms from “Biological Process”, “Molecular Function”, and “Cellular Components” from RRHO-determined “uu” (upregulated-upregulated) genes from NSD1 cKO and DNMT3A^KO/+^ mouse cortex using clusterProfiler (Wu et al., 2021; Yu et al., 2012). Data are from NSD1 cKO and control mouse whole cortex at two or eight weeks. Per time point: 2-weeks, n = 5 bioreplicates for H3K36me2, H3K36me3, DNMT3A ChIP-Rx. 8-weeks, n = 3 bioreplicates for DNA methylation, n = 6 bioreplicates for total RNA-seq. TADs were derived from Hi-C analysis of cerebral cortex (Clemens et al., 2019; Dixon et al., 2012). *p < 0.05, **p < 0.01, #p < 10^-5^, †p < 10^-10^, §p < 10^-15^.

In addition to genome-wide deposition of mCA, DNMT3A also deposits mCG at kilobase-scale regions across the neuronal genome during postnatal maturation of the brain (Christian et al., 2020; Lister et al., 2013). Notably, heterozygous disruption of DNMT3A in a mouse model of TBRS leads to loss of mCG at differentially methylated regions (DMRs) that overlap with these sites (Christian et al., 2020). We therefore sought to assess the impact of NSD1 loss on mCG levels at discrete regions across the genome and interrogate if these sites overlap with regions disrupted in models of DNMT3A disorders. To comprehensively assess how mCG is affected upon reduction of NSD1 activity, we performed BSmooth analysis to identify mCG-DMRs *de novo* in the NSD1 cKO (Hansen et al., 2012). Our analysis of mCG identified 3295 hyper-DMRs and 7447 hypo-DMRs (Figure 6D). The identification of a preponderance of hypo-DMRs further supports a role for H3K36me2 in targeting deposition of DNA methylation in the neuronal genome. Analysis of the location of these DMRs showed that loss of NSD1 leads to changes in mCG at important regulatory sites including genes, promoters, CpG shores, CpG islands, and enhancers (Figure 6E,F). Notably, we found that adult DMRs, regions of the genome that acquire mCG during postnatal development in a DNMT3A-dependent manner (Lister et al., 2013), were most enriched for overlap with NSD1 cKO DMRs (Figure 6E). These sites often overlap regulatory elements that undergo repression during postnatal development. To further analyze the effects of NSD1 deletion on mCG across these important genomic elements we expanded our analysis to quantitatively assess effects on CG sites across the classes of sequences genome-wide. Analysis of changes in H3K36me2, DNMT3A, and DNA methylation levels across these regulatory sites revealed concordant reduction of H3K36me2 and DNMT3A binding, with loss of mCA and mCG in adult DMRs exhibiting the largest reductions in methylation (Figure 6F).

Notably, adult DMRs have been shown to be susceptible to CG hypo-methylation in DNMT3A heterozygous mutant (DNMT3A^KO/+^) mice, suggesting similar losses of mCG between TBRS models and NSD1 cKO mice. To more directly assess if the effects on mCG are similar between NSD1 and DNMT3A mutant models we determined the degree of overlap between DMRs called in the NSD1 cKO versus in the DNMT3A^KO/+^ and found significant overlap between hypo-DMRs called in each model (Figure 6I). These results indicate that specific mCG sites are sensitive to loss of NSD1 and overlap with aberrantly methylated sites in DNMT3A models, implicating common DMRs across these disease models that may affect gene expression. Our findings further support a model in which NSD1-mediated H3K36me2 deposition functions as a direct upstream regulator of DNMT3A to put down large-scale mCA and site-specific mCG in the postnatal brain.

We next investigated whether the epigenetic changes we observe in the NSD1 cKO results in altered gene expression in the brain. We performed differential gene expression analysis by total RNA-seq of eight week control and NSD1 cKO whole cortex, identifying 217 (177 up, 40 down) significantly dysregulated genes (FDR<0.1) (Figure 6G). Analysis of changes in H3K36me2, H3K36me3, DNMT3A, and DNA methylation revealed reduced H3K36me2, DNMT3A, and mCA levels and increased H3K36me3 levels at aberrantly upregulated genes (Figure 6H), supporting a role for epigenetic dysregulation in driving these effects. Analysis of levels of these chromatin and DNA modifications for the TADs in which the significantly called dysregulated genes were found (Figure S6H) revealed similar changes in these modifications at the TAD-level. In addition, we detected overlap between significantly dysregulated genes and TADs with significant changes in H3K36me2 (Figure S6G). Together, these results suggest that the gene dysregulation observed in the NSD1 cKO is due in part to changes in H3K36me2 and subsequent mCA levels at these genes within affected TADs.

In light of the overlapping epigenetic disruption we detect in NSD1 and DNMT3A mutant mice, and the fact that neurodevelopmental disorders caused by mutation of these genes share considerable clinical phenotypic overlap (Tatton-Brown et al., 2017), we investigated if shared gene expression effects may occur across mutant mouse models. Identification of common dysregulated genes in these mice would provide candidates for drivers of shared pathology in disease. We therefore examined the extent of overlap in gene dysregulation between NSD1 cKO and DNMT3A mutant models. Both models display a small number of genes that reach statistical significance for dysregulation. Despite this fact, overlap analysis of significantly dysregulated genes in the NSD1 cKO (217 genes) and the DNMT3A^KO/+^ (65 genes) revealed significant enrichment of co-upregulated genes across the two mouse models (Figure S6I).

Analysis of multiple models of NDD has demonstrated that gene expression effects in the brain caused by these mutations result in widespread but subtle dysregulation of many genes that is not captured by analysis of statistically significant genes alone (Christian et al., 2020; Clemens and Gabel, 2020; Nord and West, 2020). Therefore, to further assess transcriptome-wide overlap in gene dysregulation between these two genotypes, we performed Rank-Rank Hypergeometric Overlap (RRHO) analysis (Cahill et al., 2018; Plaisier et al., 2010) of gene expression changes in the NSD1 cKO and DNMT3A mutant models. RRHO revealed widespread and significant concordance for upregulated genes between the NSD1 cKO and DNMT3A mutant models, indicating global similarity between gene de-repression effects caused by disruption of these genes (Figure 6J, S6J). Such broad, shared de-repression is consistent with the loss of the repressive mCA mark across the genome, and points to substantial overlap in gene dysregulation in these models.

Having detected overlap in epigenetic and gene expression changes in the NSD1 cKO and DNMT3A mutant models, we sought to investigate the functional pathways impacted by these shared effects. We therefore performed Gene Ontology analysis (Wu et al., 2021; Yu et al., 2012) of concordantly dysregulated genes identified by RRHO. This analysis detected terms associated with key neurodevelopmental processes during postnatal brain development, including synapse organization and axon guidance (Figure 6K, S6K). Identification of terms such as glutamatergic synapse and ion channel activity further suggest disrupted neurotransmission manifesting due to transcriptional dysregulation upon DNMT3A and NSD1 mutation. Thus, the role of NSD1 as a direct upstream regulator of DNMT3A binding and mCA deposition can influence critical pathways involved in developing and mature neural circuits. Overall, these findings identify key functional pathways in the brain with shared disruption upon NSD1 or DNMT3A loss and provide evidence of etiological convergence at the molecular level for NDDs caused by mutation of these genes.

## Discussion

In this study we have interrogated the mechanisms of DNMT3A targeting in the brain to uncover a major role of H3K36me2 in dictating neuronal mCA accumulation. Our findings are consistent with a model in which the activity of NSD1 is directed by genome topology to broadly pattern varying levels of H3K36me2 across the neuronal genome in the developing nervous system. This regional H3K36me2 can then be modified through depletion at genes when H3K36me2 is converted to H3K36me3 by active transcription. During the postnatal period DNMT3A is recruited to H3K36me2 sites through its PWWP domain to establish mCA patterns. mCA patterns are then read out by MeCP2 and possibly other proteins to repress regulatory elements and tune transcription of essential gene programs in the mature brain. Mutation of NSD1, DNMT3A, or MeCP2 can disrupt the establishment or read out of this critical H3K36me2-mCA pathway, contributing to nervous system dysfunction in human NDDs.

Our study also demonstrates that deposition of H3K4me2/3 at active regulatory sites is associated with further blocking of DNMT3A activity, presumably by abrogating the capacity of the ADD domain to activate the enzyme and driving focal depletion of mCA. This blocking of mCA is consistent with regulation of DNMT3A described for mCG deposition in non-neural systems and is an important additional determinant of the levels of mCA at enhancers and promoters. Thus, H3K4 methylation in the early postnatal period can also impact subsequent mCA-directed repression of enhancers and gene expression in the mature brain. Notably, disruption of both H3K4 methyltransferases (KMT2A, KMT2C, KMT2D) and demethylases (LSD1, KDM5C) have been observed in individuals with NDD, suggesting additional disorders in which the neuronal DNA methylome may be impacted (Faundes et al., 2018; Iwase et al., 2007; Ng et al., 2010; Pilotto et al., 2016; Vallianatos and Iwase, 2015; Yuen et al., 2017). Future studies can dissect the role of H3K4 methylation in patterning mCA and examine if this process contributes to these NDDs.

By integrating analyses of gene expression and regional methylation together and exploring underlying chromatin marks associated with these phenomena, our study reconciles two distinct mechanisms of mCA patterning that had been independently described (Clemens et al., 2019; Stroud et al., 2017). Rather than working through completely independent mechanisms or interacting through complex non-linear processes, however, we find that regional set-points and gene-expression-associated depletion of mCA can be explained by two steps in H3K36me2 patterning: regional deposition of the mark followed by selective conversion to H3K36me3. Thus, our findings provide a unifying framework to explain how neurons acquire their unique methylomes.

Our findings suggest that reduction of H3K36me2, perhaps due to conversion of H3K36me2 to H3K36me3, in highly expressed genes is associated with reduced mCA accumulation at these regions. In agreement with this model, biochemical studies comparing PWWP domain binding of DNMT3A to these two marks support a higher affinity for H3K36me2 over H3K36me3 (Dukatz et al., 2019; Weinberg et al., 2019). However, DNMT3A displays stronger affinity to H3K36me3 compared to unmodified histones in several studies (Dukatz et al., 2019; Weinberg et al., 2019; Xu et al., 2020), and thus it remains unclear if this additional affinity plays a role in DNMT3A targeting. Previously, steric hinderance due to passthrough of the transcriptional machinery has been invoked as a possible mechanism by which DNMT3A may be excluded from active genes (Stroud et al., 2017), and it remains possible that, in addition to a reduced affinity upon conversion of H3K36me2 to H3K36me3, steric effects contribute to loss of DNMT3A binding to intermediate affinity H3K36me3 sites within genes. Future studies that modulate H3K36me3 or transcription through genes directly will be necessary to further dissect this process.

Our integrated analysis of chromatin marks with DNMT3A binding and mCA deposition across gene regulatory elements establishes H3K36me2 as a strong predictor of DNMT3A-mediated mCA deposition genome-wide. Histone variant H2Az also displayed a strong negative relationship with DNMT3A binding and mCA deposition at promoters (Figure 4A), which is supported by previous studies with similar results between H2Az and mCG (Edwards et al., 2010). Interestingly, our random forest algorithm also uncovered H3K27 and H3K9 methylation as predictors of DNMT3A-mediated mCA deposition (Figure 4C,D). While previous studies have detected a positive association between H3K9 methylation and DNA methylation (Epsztejn-Litman et al., 2008; Meissner et al., 2008), the interplay between H3K27 methylation and DNA methylation is complex and remains incompletely defined. However recent studies have suggested that H3K36me2 may coordinate with H3K27 methylation and other chromatin marks to dynamically regulate DNMT3A binding and DNA methylation patterning (Chen et al., 2022; Gu et al., 2022; Streubel et al., 2018). Future studies investigating the contribution of these other chromatin-associated marks will be important to comprehensively understand the complex role and predictive capacity of H3K36me2 on DNMT3A-mediated mCA patterning in the brain and in neurodevelopmental disease.

Our analysis in cerebral cortex tissue establishes a model for how targeting of mCA is carried out during postnatal development. Notably however, mCA has been shown to exhibit highly cell-type specific patterns across different types of neurons in the brain (Liu et al., 2021; Luo et al., 2017; Mo et al., 2015). Thus, the patterns we observe here reflect the averages across many cell types. While the mechanisms we uncover provide conceptual insight into how DNMT3A is targeted, these mechanisms are likely to be differentially utilized in cell types to create unique mCA patterns. For example, differential activation of genes can drive expression-associated protection from mCA at different gene sets across cell types (Stroud et al., 2017, 2020). It will be important to explore how H3K36me2, and regional set-point methylation contribute to shared and distinct mCA patterns across neuronal cell types.

TBRS and Sotos syndrome are marked by multiple overlapping phenotypes including overgrowth, intellectual disability, and a high penetrance of autism. Our findings suggest that disrupted deposition of mCA in broad domains and loss of discrete regions of mCG targeting upon NSD1 or DNMT3A mutation may contribute to shared cognitive and behavioral phenotypes across these disorders. Our gene expression analysis indicates that these overlapping epigenetic effects in NSD1 and DNMT3A mutant mice drive concordant gene dysregulation that are likely to drive these nervous system deficits. Overgrowth in these disorders is likely due to epigenetic alterations in dividing cells during early development. The convergence of epigenetic disruption across these disorders both inside and outside the nervous system may present targets for shared therapeutic development. Notably, in addition to specific megabase-scale domains being affected by NSD1 deletion in the brain, there is a trend toward reduced global mCA upon NSD1 mutation that mimics global reductions observed in TBRS models (Figure S6E). Thus, augmenting existing DNMT3A activity in neurons with genetic approaches or enzyme agonists may ameliorate epigenetic anomalies in both disorders, and may represent a feasible strategy for shared therapeutic development.

Notably while knockdown or conditional deletion of NSD1 alters broad H3K36me2 patterns and mCA in neurons, loss of these marks is incomplete in this mutant. Partial disruption of H3K36me2 could be due, in part, to the acute nature of NSD1 inactivation combined with the long half-life of histones and histone modifications in postmitotic neurons (Maze et al., 2015). However, it is likely that other H3K36 methyltransferases expressed in the developing nervous system, including ASH1L and NSD2, can deposit and maintain this mark. Notably, these genes are also mutated in NDD with autistic features or cognitive deficits (Bergemann et al., 2005; Sanders et al., 2015). Disruption of these modifiers may lead to both overlapping or distinct effects on H3K36me2 and DNA methylation in the brain compared to loss of NSD1 and DNMT3A. Furthermore, SETD2, the methyltransferase that converts H3K36me2 to H3K36me3 in expressed genes is also mutated in NDD (Lumish et al., 2015), raising the possibility that, in addition to direct effects of loss of H3K36me3, expression-associated mCA depletion in genes may be affected in individuals with SETD2 mutations. Thus, multiple additional genetic causes of NDD may impact the mCA pathway, and alterations in mCA may, in turn, contribute to disease pathology across these additional disorders. Future studies can examine the degree to which H3K36me2 and mCA are altered in models of these NDDs and dissect the shared and distinct consequences of loss of H3K36me2. This and future works can map the unique neuronal epigenome as a growing point of convergent disruption across autism and related NDDs.

## Supporting information

Tabel S1

## Acknowledgements

We thank members of the Gabel lab, as well as J. Edwards, N. Mosammaparast, and J. Yi, for providing vital support and feedback on the manuscript. Next-Generation Sequencing was carried out through the Genome Technology Access Center at the McDonnell Genome Institute and The Edison Family Center for Genome Sciences and Systems Biology at Washington University in St. Louis. This work was supported by NIH NICHD 1F30HD110156-01 to N.H. and NIMH R01MH117405 to H.W.G.

## Author Contributions

Conceptualization and Methodology, N.H., D.Y.W., and H.W.G.; Experimentation, N.H. A.W.C. S.A.N.; Formal Analysis, N.H., D.Y.W.; Manuscript – Writing, N.H., D.Y.W., H.W.G.; Manuscript – Review & Editing, N.H, D.Y.W., H.W.G.

## STAR Methods

### Key Resources Table

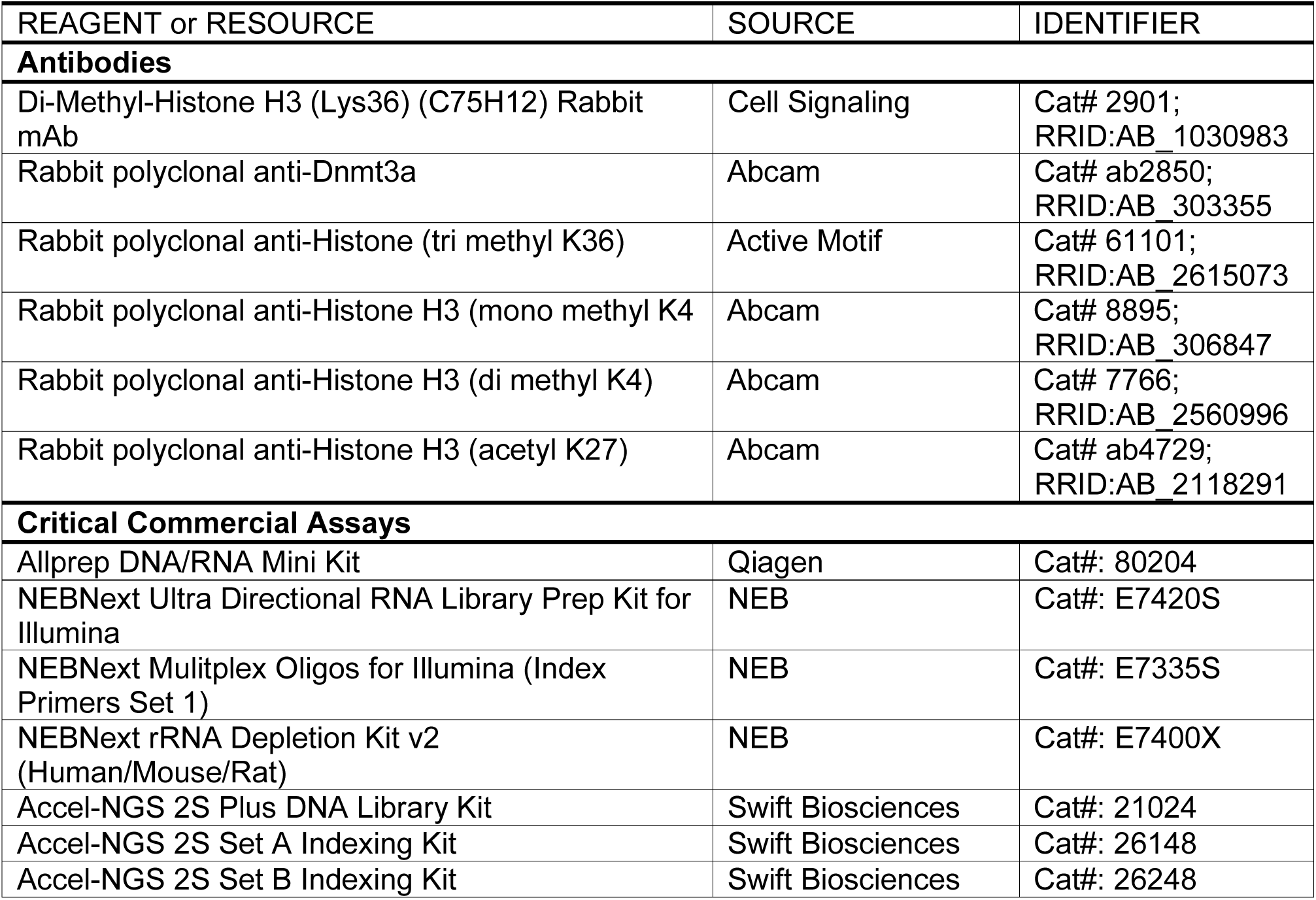
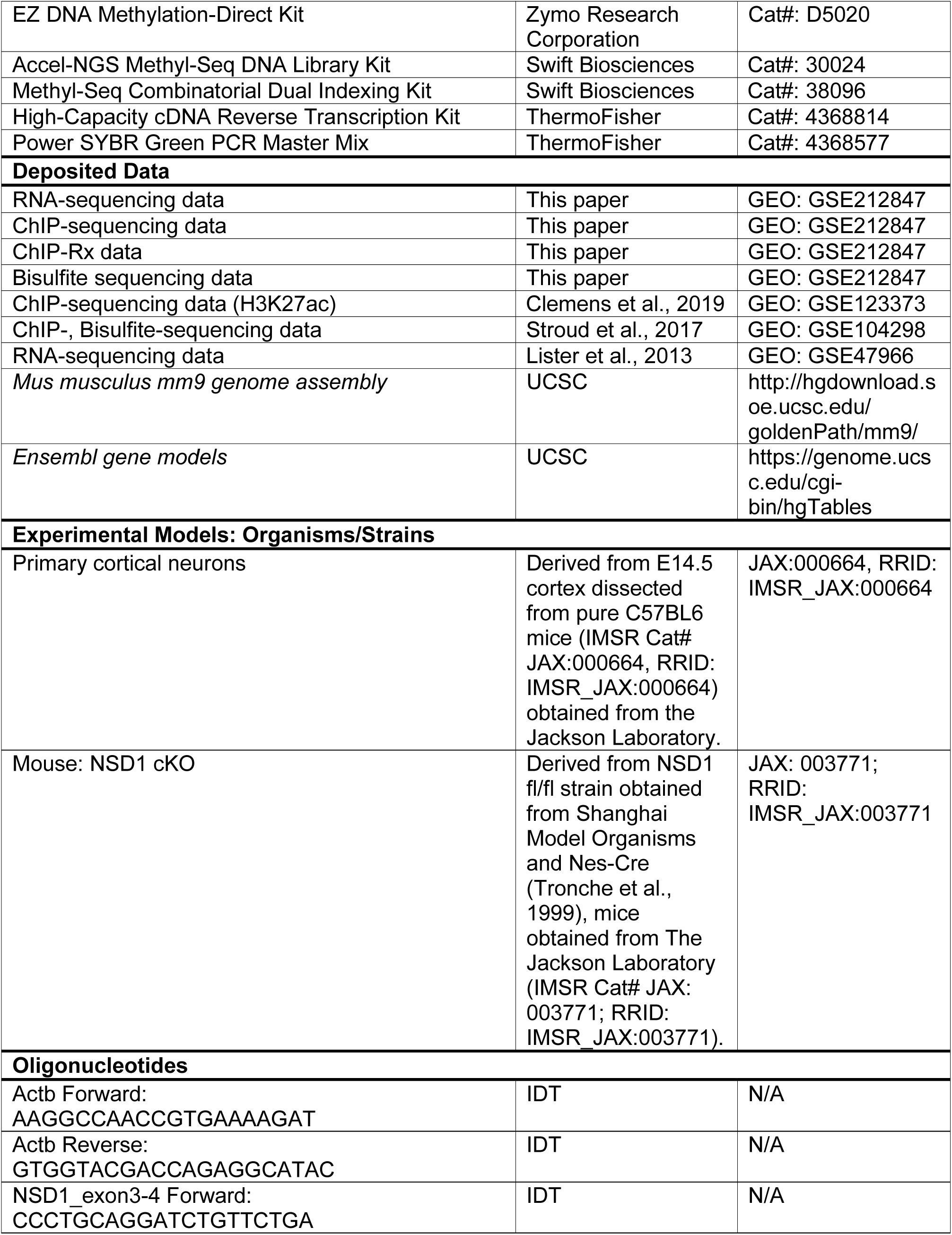
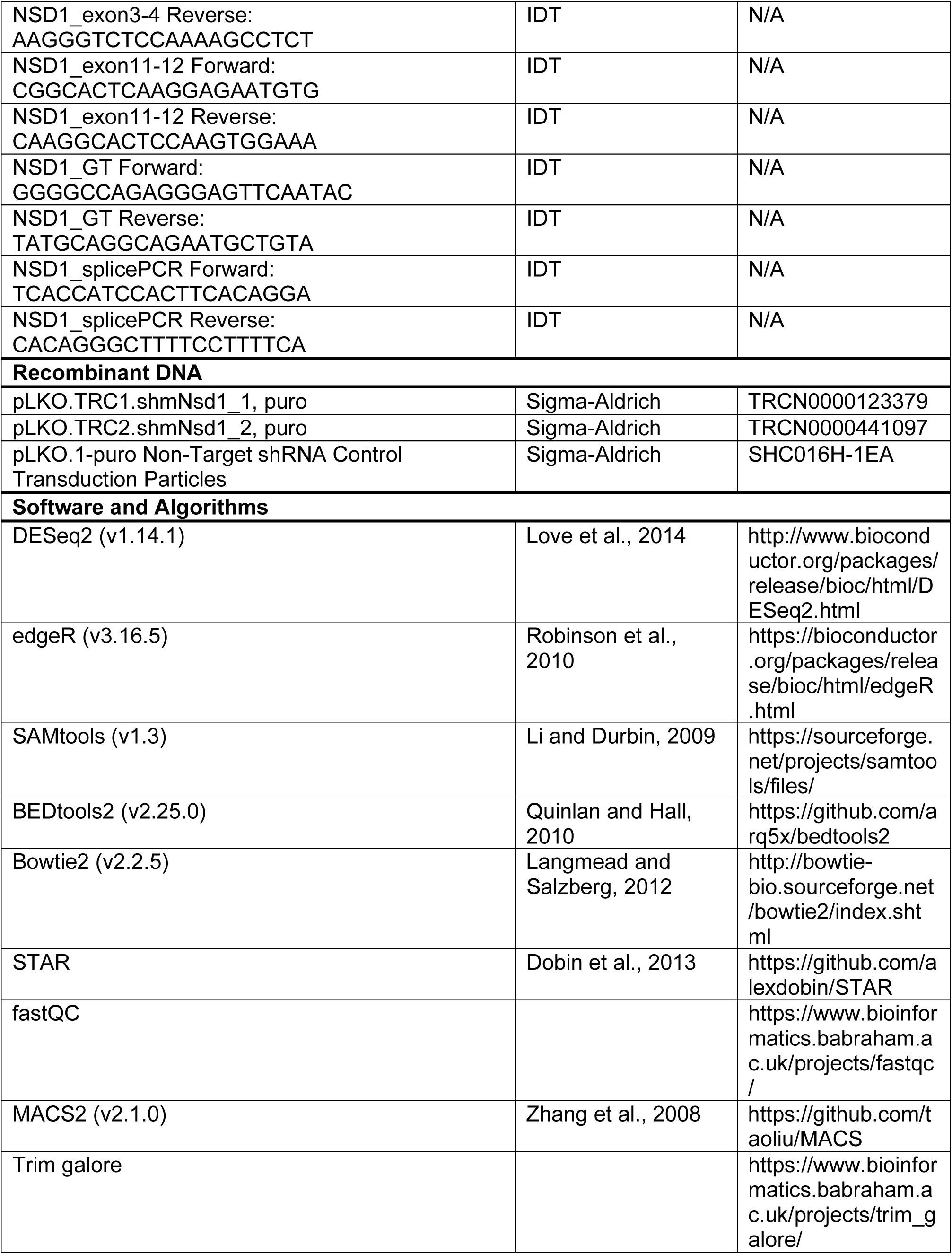
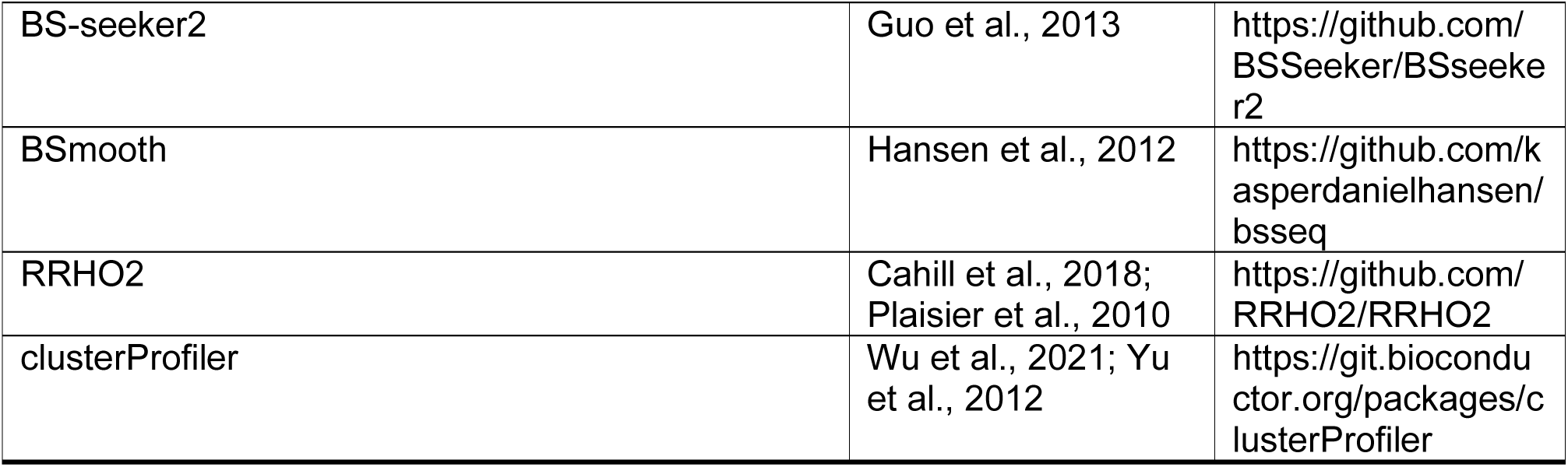

### Contact for Reagent and Resource Sharing

Requests for reagents and resources should be directed towards the Lead Contact, Harrison Gabel.

### Experimental Model and Subject Details

#### Mice

Male and female heterozygous Nsd1^flx/+^ mice were generated by Shanghai Model Organisms. Nsd1^flx/flx^ mice were generated by crossing Nsd1^flx/+^ males to Nsd1^flx/+^ females. Female Nsd1^flx/flx^ mice were crossed to male B6.Cg-Tg(Nes-cre)1Kln/J (Nestin-Cre^+/-^) to generate Nsd1^flx/+^; Nestin-Cre^+/-^. Male Nsd1^flx/+^; Nestin-Cre^+/-^ were then crossed to female Nsd1^flx/flx^ to generate Nsd1^flx/flx^ Tg(Nes-cre)1Kln/J conditional knockout mice (NSD1 Nestin-cKO). Genotypes of each mouse was confirmed using primers surrounding floxed region (F: GGGGCCAGAGGGAGTTCAATAC, R: TATGCAGGCAGAATGCTGTA). Relative abundance of NSD1 long isoforms in NSD1 control and cKO mice were assess using primers targeting exons 2 through 5 (F: TCACCATCCACTTCACAGGA, R: CACAGGGCTTTTCCTTTTCA).

### Methods

#### Total DNA/RNA isolation

Cerebral cortex was dissected on ice in phosphate buffered saline (PBS) from NSD1 Nestin-cko and wild-type litter mates at 8 weeks old. Total DNA and RNA were extracted from 300,000 primary cultured neurons or 1/10 of a whole cortex using RLT+ buffer following the Allprep DNA/RNA Mini Kit (QIAGEN).

#### qRT-PCR

RNA isolated from neuronal cortical culture or mouse cerebral cortex tissue was reverse transcribed using the High-Capacity cDNA Reverse Transcription Kit (Applied Biosystems). Nsd1 and Actb were measured by qPCR using the Power SYBR Green PCR Master Mix and primers Actb (F: AAGGCCAACCGTGAAAAGAT, R: GTGGTACGACCAGAGGCATAC) and Nsd1 (PNCs - F: CGGCACTCAAGGAGAATGTG, R: CAAGGCACTCCAAGTGGAAA; Cortex tissue - F: CCCTGCAGGATCTGTTCTGA, R: AAGGGTCTCCAAAAGCCTCT). Relative quantity of Actb to Nsd1 cDNA was determined by comparing the Ct of each primer set in each sample to a standard curve and then normalizing the NSD1 signal by the ACTB signal.

#### Chromatin immunoprecipitation from primary neuron cultures

Cross-linking ChIP in primary neuron cultures was performed using 700,000 cells per immunoprecipitation. Medium was aspirated and cells were cross-linked directly on the plate using 1% paraformaldehyde for 10 min at room temperature with gentle shaking. Glycine was added to quench (final concentration 125 mM, incubated for 5 min at room temperature), then cells were washed twice with cold PBS (with PMSF). To obtain soluble chromatin extract, cells were resuspended in 1ml L1 (50 mM HEPES, 140 mM NaCl, 1 mM EDTA, 1mM EGTA, 10% glycerol, 0.5% NP-40, 0.25% Triton X-100, 1% Na butyrate and 1× Complete protease inhibitor), scraped off the plates, and incubated on ice for 5 min with occasional mixing. Cells were centrifuged and resuspended again in L1. Samples were centrifuged and resuspended in 1ml L2 (10 mM Tris-HCl pH 8.0, 200 mM NaCl, 1% Na butyrate, 1% Triton X-100, 0.5% SDS, and 1× Compete protease inhibitor). Finally, samples were centrifuged and resuspended in 300µl L3 (10 mM Tris-HCl pH 8.0, 1 mM EDTA, 1 mM EGTA, 1% Na butyrate, and 1× Complete protease inhibitor). Chromatin extracts were then sonicated for 30 min (30 cycles: 30s on, 30s off) using a Diagenode Bioruptor sonicator. The lysates were incubated with 15 μl protein A Dynabeads (Invitrogen) bound to anti-H3K36me2 (0.1µg, Cell Signaling 2901) or anti-DNMT3A (4µg, Abcam ab2850) and incubated overnight at 4 °C with 20µl kept as input DNA. Magnetic beads were sequentially washed with low-salt buffer (150 mM NaCl; 0.1% SDS; 1% Triton X-100; 2 mM EDTA and 20 mM Tris-HCl), high salt buffer (500 mM NaCl; 0.1% SDS; 1% Triton X-100; 2 mM EDTA and 20 mM Tris-HCl), LiCl buffer (150 mM LiCl; 0.5% Na deoxycholate; 1% NP-40; 1 mM EDTA and 10 mM Tris-HCl) and TE buffer (1 mM EDTA and 10 mM Tris-HCl). Beads were eluted in TE + 1%SDS and incubated for 30 min at 65 °C for two rounds. The eluate was reverse cross-linked overnight at 65 °C. The eluate was then treated with RNase A for 1 h at 37 °C and with Proteinase K (Roche) for 2 h at 55 °C and DNA was recovered using a Qiagen PCR purification kit.

#### Chromatin immunoprecipitation from tissue

Chromatin immunoprecipitation was performed as previously described (Clemens et al., 2019; Cohen et al., 2011). Cerebral cortex was dissected on ice in PBS from NSD1 Nestin-cKO and control littermates at 2- and 8-weeks old (2-weeks: n = # pairs, # male, # female, 8weeks: n = # pairs, # male, # female). The tissue was flash-frozen in liquid nitrogen and stored at −80°C. Chromatin was fragmented with the Covaris S220 sonicator (5% Duty Factory, 140Peak Incidence Power, 200 cycles per burst, milliTUBE 1mL AFA Fiber). After centrifugation, samples were spiked with soluble chromatin from Drosophila S2 cells to comprise 5% of total chromatin in the lysate. ChIP was performed with H3K36me2 antibody (0.1 µg, Cell Signaling 2901), H3K36me3 antibody (0.2 µg, Active motif pAb), H3K27me2 antibody (0.2 µg, Cell Signaling 9728), H3K27me3 antibody (0.5 µg, Active motif pAb), H3K27ac antibody (0.1µg; Abcam ab4729), and DNMT3A antibody (4 µg, Abcam ab2850). Libraries were generated using Accel-NGS 2S Plus DNA Library Kit (Swift Biosciences). Libraries were pooled to a final concentration of 10nM and sequenced using Illumina HiSeq 3000 with the Genome Technology Access Center at Washington University in St. Louis, typically yielding 15-40 million single-end reads per sample.

#### Whole genome bisulfite sequencing

DNA was isolated from primary neuron cultures or mouse brain tissue using the Allprep DNA/RNA Mini Kit (QIAGEN). Libraries were prepared for sequencing using the Accel-NGS Methyl-Seq DNA Library Kit (Swift, 30024) and the EZ DNA Methylation-Direct Kit (Zymo, D5020) was used for bisulfite conversion. For these samples, 50 ng of DNA was fragmented for 45 s with the Covaris S220 sonicator (10% Duty Factory, 175 Peak Incidence Power, 200 cycles per burst, microTUBE 200µL AFA Fiber). DNA was then purified using 0.7 volumes of SPRISelect Beads (Beckman Coulter Life Sciences) to select for long DNA inserts for sequencing. Samples underwent bisulfite conversion under the following cycling conditions: 98°C, 8 min; 64°C, 4.25 h, 4°C hold. Libraries were PCR-amplified for 10 cycles. Libraries were then pooled to a final concentration of 5nM and sequenced using Illumina HiSeq 3000 with the Genome Technology Access Center at Washington University in St. Louis, typically yielding 100 million paired-end reads per sample.

### Quantification and statistical analysis

#### Machine Learning

Random forest models were built with python sklearn with the parameters n_estimators=100, min_samples_split=2, criterion=’mse’. 5-fold cross-validation was used to evaluate performance. Individual models were built and assessed for each set of regions, using input-normalized values for the ChIP-signal in each region. Region orders were randomized before cross-validation, using random.seed(9115).

Feature importances of recursive feature elimination were plotted as a series of stacked barplots, with their heights scaled to the accuracy of the model with the given number of features removed. Linear models were built with R using input-normalized data.

#### Whole-genome bisulfite analysis

Bisulfite sequencing analysis was performed as previously described (Clemens et al., 2019), with the addition of nonconversion correction for regional %mC assessment. Briefly, data were adaptor-trimmed, mapped to mm9, then deduplicated and called for methylation using BS- seeker2. Methylation levels across regions were assessed using bedtools map -o sum, summing the number of reads mapping to Cs (supporting mC) and the number of reads mapping to Cs + Ts (supporting C) in the region, then dividing those two numbers (Quinlan and Hall, 2010). The coverage from split sequencing runs were pooled together before assessing %mC, while the %mC values from biological replicates were averaged together.

Regions were adjusted for nonconversion rate as measured by % methylation in Lambda spike-ins per-sample, as in (Lister, 2013). Briefly, following %mC calculation, the lambda % methylation value was subtracted from the calculated %mC value. If the corrected value was below 0, the %mC value was brought up to 0.

#### RNA sequencing analysis

RNA sequencing analysis was performed as previously described (Clemens et al., 2019). Briefly, FASTQ files were adapter and quality-trimmed with Trim Galore and filtered for rRNA sequences with Bowtie. The remaining reads were aligned to the mm9 genome with STAR (Dobin et al., 2013). Reads mapping to multiple regions were removed, and the remaining uniquely mapping reads were converted to BED files and separated by intronic and exonic reads. These BED files were then used to assess coverage in genes using bedtools coverage -counts.

DESeq2 was used to identify significant differentially expressed genes (FDR<0.1) in the control versus NSD1 cKO cerebral cortex.

#### Chromatin immunoprecipitation analysis

ChIP sequencing analysis was performed as previously described (Clemens et al., 2019). Briefly, reads were mapped to mm9 with bowtie2, then deduplicated with picardtools MarkDuplicates. bedtools coverage -counts was used to assess ChIP signal at the various genomic regions examined.

The global change in ChIP enrichment was calculated using ChIP-Rx, using an exogenously added Drosophila chromatin (5% of sample) as an internal control (Orlando et al., 2014). For each ChIP-seq sample, the ChIP-Rx ratio was calculated as follows:

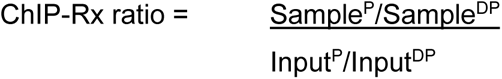

in which Sample^P^ and Sample^DP^ refer to the percentages of reads that aligned to the mouse (mm9) and Drosophila (dm6) genome, respectively (likewise, Input^P^ and Input^DP^ refer to the percentage of input that aligned to each genome). Given that only H3K36me2 ChIP-seq had a robust change in global enrichment (Figure S6E), the H3K36me2 ChIP-seq track was normalized using bedtools genomecov -scale with the scale value equal to the percent global change in H3K36me2 enrichment in the NSD1 cKO per bioreplicate.

### Controlled resampling

A similar resampling approach was used as previously described (Clemens et al., 2019). Briefly, genes were sorted based on some criteria (e.g. expression). Then, for every entry in a set of interest (e.g. MeCP2-repressed genes), an entry from a control set (e.g. all other genes) with a similar rank as the entry of interest was taken, generating a control set of the same size and variable distribution as the set of interest.

### TAD cross-correlation

Cross-correlation matrices (e.g., Figure 5E, Figure S2A) were generated as previously described (Clemens et al., 2019). Briefly, each TAD was divided into 10 equally-sized bins. 10 identically sized bins were subsequently appended up and down-stream of the TAD, resulting in a single matrix of 30 bins. Each column of this matrix was then correlated against each other, making a 30 x 30 correlation matrix, which was plotted in heatmap form.

To generate heatmaps of correlations of fold-changes of H3K36me2, DNMT3A, and mCA of regions inside versus outside of TADs (Figure 5E) as well as H3K36me2 levels in and around all TADs (Figure S2A), each element was paired to each other element on the same chromosome. Each pair was then assessed if they paired within or between TADs. Because H3K36me2, DNMT3A, and mCA vary with genomic distance, each intra-TAD pair was matched to the inter-TAD pair with the most similar distance between elements. Spearman correlations were then calculated on the two distance-matched sets.

### Identification of NSD1 changed regions

Regions (10kb bins across the genome, TADs) were assessed for ChIP signal in control and NSD1 knockdown or NSD1 cKO samples. edgeR was used to identify regions with significantly changed enrichment of H3K36me2 (FDR <0.1).

For Figure 6B, edger counts were normalized by the percent change observed in the NSD1 cKO by ChIP-Rx to obtain the x-axis range that reflects this global shift in enrichment (DGEList() object$samples$norm.factors * ChIP-Rx calculated percent change). However, to obtain the genomic regions with the most significant change in H3K36me2 for subsequent analyses, we used non ChIP-Rx normalized edgeR outputs.

### Tiling of genome into 5kb and 10kb bins

Genome was divided into 5kb and 10kb bins with a method similar to (Stroud et al., 2017). Briefly, bins were placed throughout the genome and labeled in a gene-centric manner – using a sorted annotation file of genes, bins were iteratively placed at the center of a gene’s TSS, then downstream of the TSS to the TES if the gene was long enough to allow at least one bin inside it. Bins were then placed immediately downstream of the gene’s TES until the bins ran into the TSS of another gene, at which point the process repeats itself until the chromosome ends.

### Rank-rank hypergeometric overlap (RRHO) analysis

For each genotype, an input gene list was created, such that the first column contained the gene name and the second column contained the gene score. The gene score was calculated as - log10(pvalue) * sign(log2FC) using values obtained from the DESeq2 output for each gene. RRHO2_initialize() was used to generate RRHO object. RRHO2_heatmap() was used to generate a heatmap of overlapping genes between two genotypes.

### Over-representation analysis with clusterProfiler

A list of genes with overlapping effects (“dd” = gene in both genotypes is downregulated, “uu” gene in both genotypes is upregulated) were extracted from the RRHO object generated using RRHO2_initialize(). enrichGO() with parameters pAdjustMethod = “fdr”, ont = “ALL”, and default minGSSize and maxGSSize was used to obtain significant GO terms in each category (“BP”, “MF”, “CC”). dotplot() with parameters showCategory=5 and split = “ONTOLOGY” was used to visualize the top five significantly enriched GO terms in each category.

### Data and Software Availability

All genomic data generated in this study have been uploaded to the NCBI GEO archive GSE212847.

## Supplemental Information

Table S1. Table of *in vitro* primary neuron culture edgeR ChIP signal outputs at each genomic region.

Table S2. Table of *in vitro* primary neuron culture DNA methylation levels at each genomic region.

Table S3. Table of *in vivo* NSD1 control and cKO cerebral cortex edgeR ChIP signal outputs at each genomic region.

Table S4. Table of *in vivo* NSD1 control and cKO cerebral cortex DNA methylation levels at each genomic region.

Table S5. Table of *in vivo* NSD1 control and cKO cerebral cortex differential gene expression and DESeq2 outputs.

## Supplemental Figure Legends

**Figure S1.**
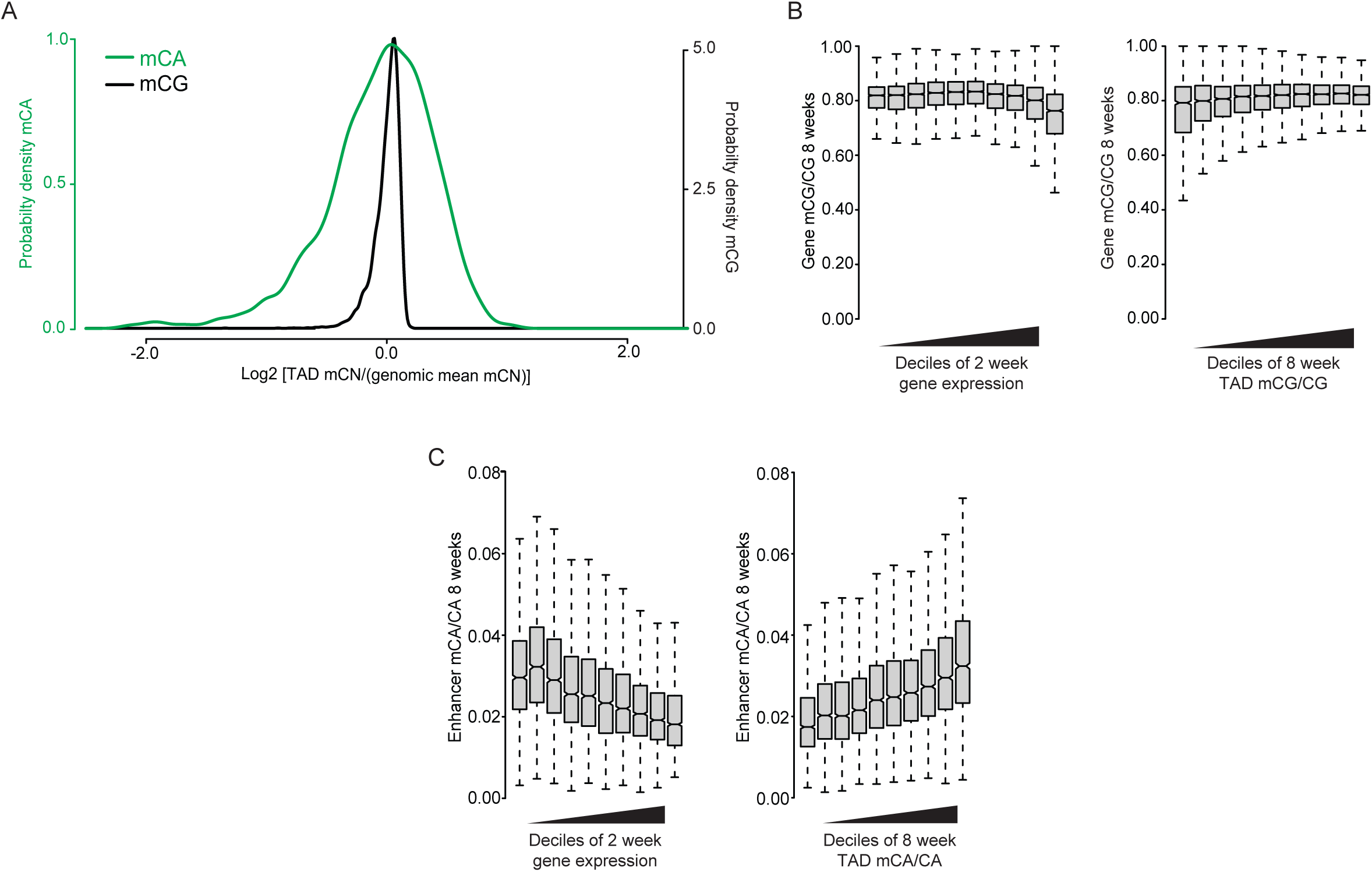
Analysis of neuronal mCA and mCG profiles associated with genomic regions and gene expression. Related to Figure 1. (A) Probability Density Functions illustrating the relative fluctuations compared to the genomic mean for levels of mCA (green) or mCG (black) in all TADs genome-wide. (B) Boxplots of mCG levels in gene bodies across deciles of gene expression (left) and TAD mCG (right). mCG levels show much more limited fluctuations associated with these two signals compared to mCA (e.g. compare to Figure 1D). (C) Boxplots of mCA levels at intragenic enhancers across deciles of gene expression (left) and TAD mCA (right). Data are from wildtype cerebral cortex at 2- and 8-weeks. 2-week RNA-seq data obtained from Lister et al., 2013. 8-week DNA methylation data obtained from Stroud et al., 2017.

**Figure S2.**
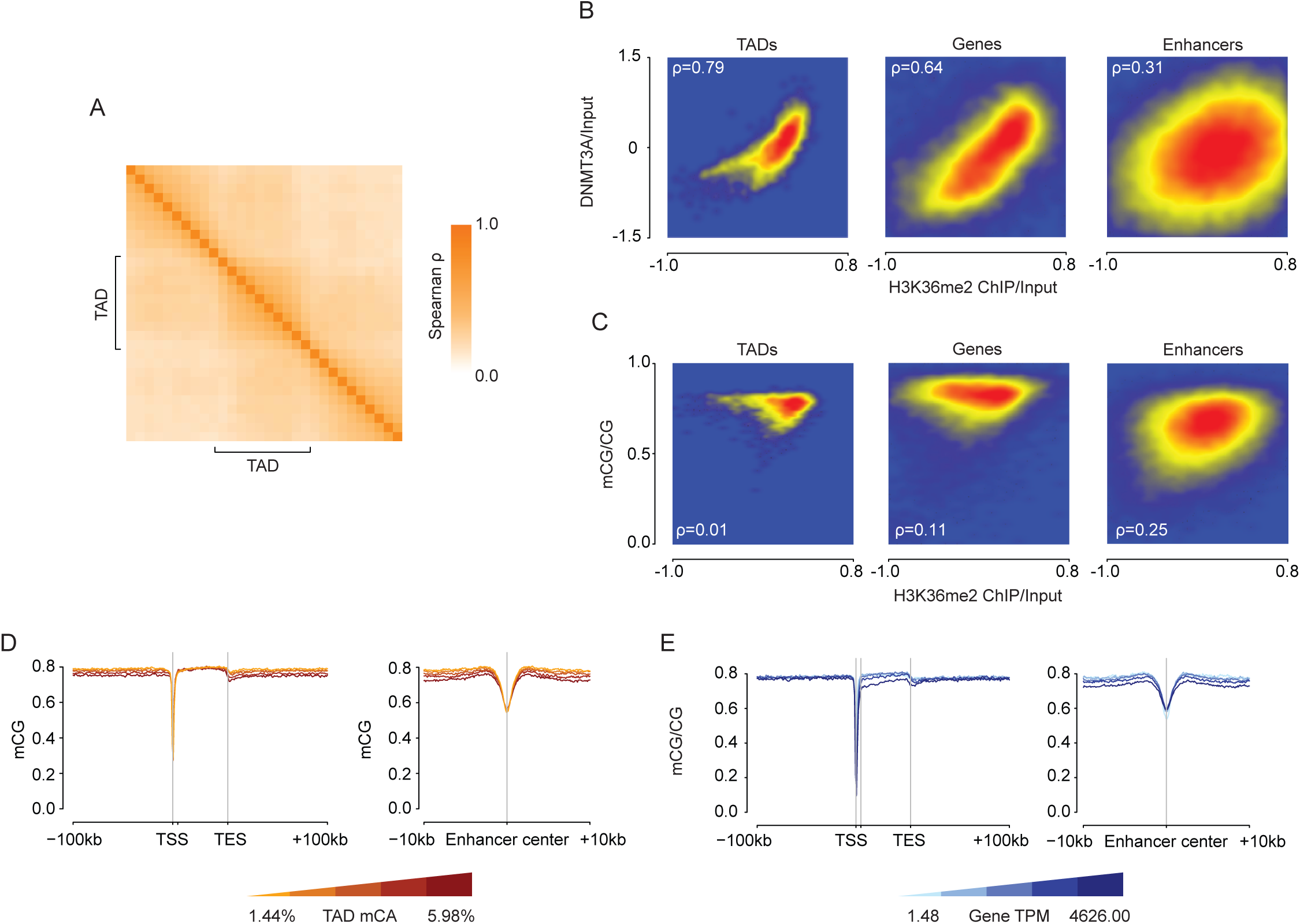
Analysis of H3K36me2 profiles and deposition of neuronal DNA methylation. Related to Figure 2. (A) Cross-correlation analysis (see *methods*) of H3K36me2 in and around all TADs in wildtype cortex. The H3K36me2 ChIP signal was assessed for 10 genomic windows within each TAD as well as for 10 equally sized windows on either side of each TAD. The correlation between the ChIP signal in these windows was assessed for all TADs genome-wide and plotted based on position relative to the TAD (see *methods*). H3K36me2 signal shows a higher correlation within TADs, compared to across TAD boundaries. (B) Comparison of H3K36me2 and DNMT3A ChIP signal at TADs, genes, and enhancers in wildtype cortex. (C) Comparison of H3K36me2 ChIP/Input and mCG/CG at TADs, genes, and enhancers in wildtype cortex. (D) Aggregate mCG levels at genes and enhancers across quintiles of TAD mCA levels. (E) Aggregate mCG levels at genes and enhancers across quintiles of gene expression. Data obtained from wildtype cerebral cortex at 2- and 8-weeks. 2-week, n = 2-3 bioreplicates for H3K36me2 and DNMT3A ChIP-seq. 2-week RNA-seq data obtained from Lister et al., 2013. 8-week DNA methylation data obtained from Stroud et al., 2017.

**Figure S3.**
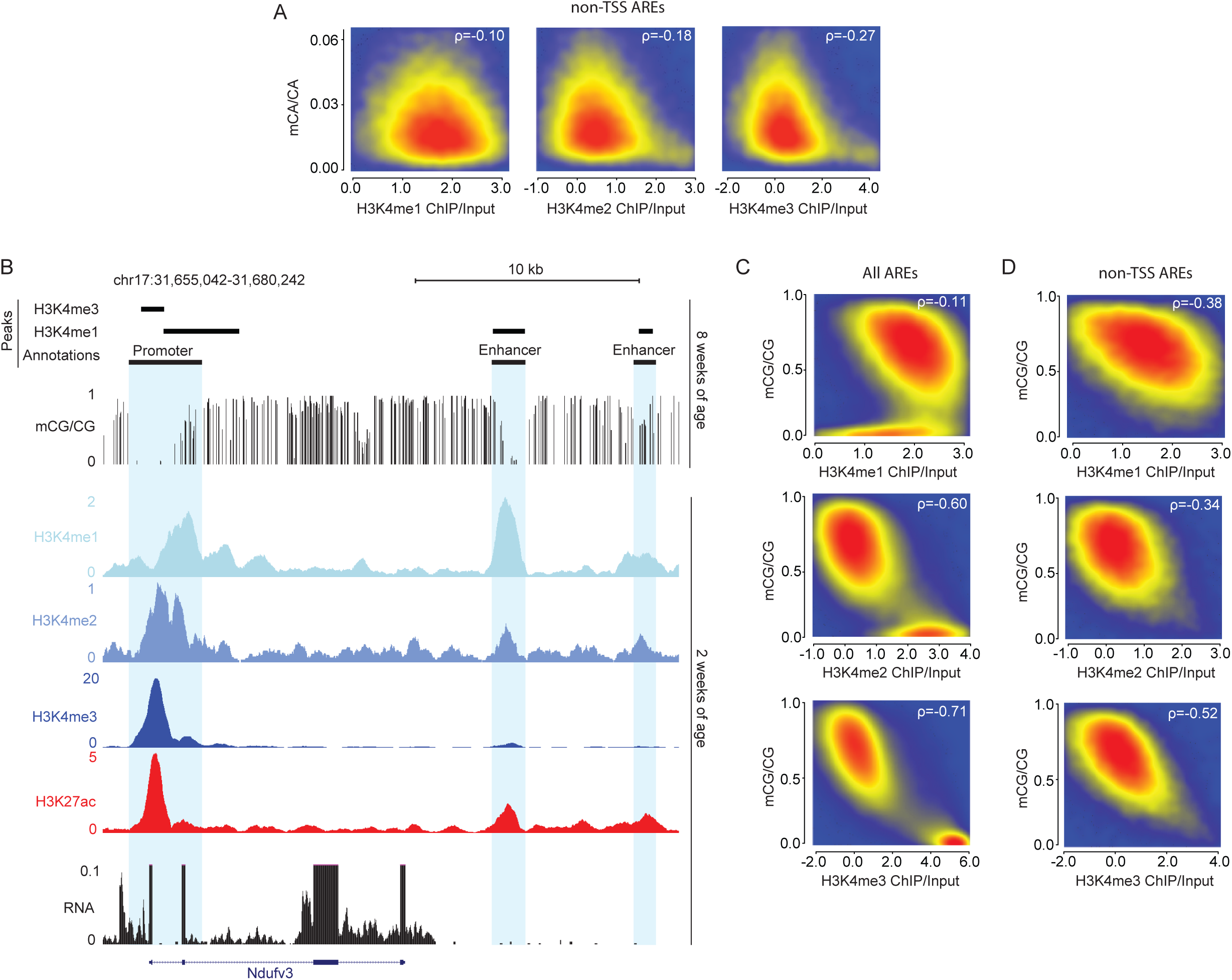
Analysis of H3K4 methylation profiles and neuronal DNA methylation. Related to Figure 3. (A) Smoothscatter plots showing relative correlation of mCA with H3K4 methylation marks at non-TSS active regulatory elements (AREs) across the genome (see *methods*). (B) Genome browser view from Figure 3 illustrating local enrichment of H3K4 methylation at active regulatory elements surrounding the *Nudfv3* gene and relative depletion of mCG at these sites. (Note: identical ChIP signals from Figure 3 are replotted here to allow comparison to mCG). (C) Smoothscatter plots showing relative correlation of mCG with H3K4 methylation marks at all active regulatory elements across the genome (see *methods*). (D) Smoothscatter plots showing relative correlation of mCG with H3K4 methylation marks at non-TSS active regulatory elements across the genome (see *methods*). Data are from wildtype cerebral cortex at 2- and 8-weeks. 2-week, n = 2 bioreplicates for DNMT3A, H3K4me1, and H3K4me2 ChIP-seq. RNA-seq data obtained from Lister et al., 2013. DNA methylation and H3K4me3 ChIP-seq data obtained from Stroud et al., 2017. H3K27ac ChIP-seq data obtained from Clemens et al., 2019.

**Figure S4.**
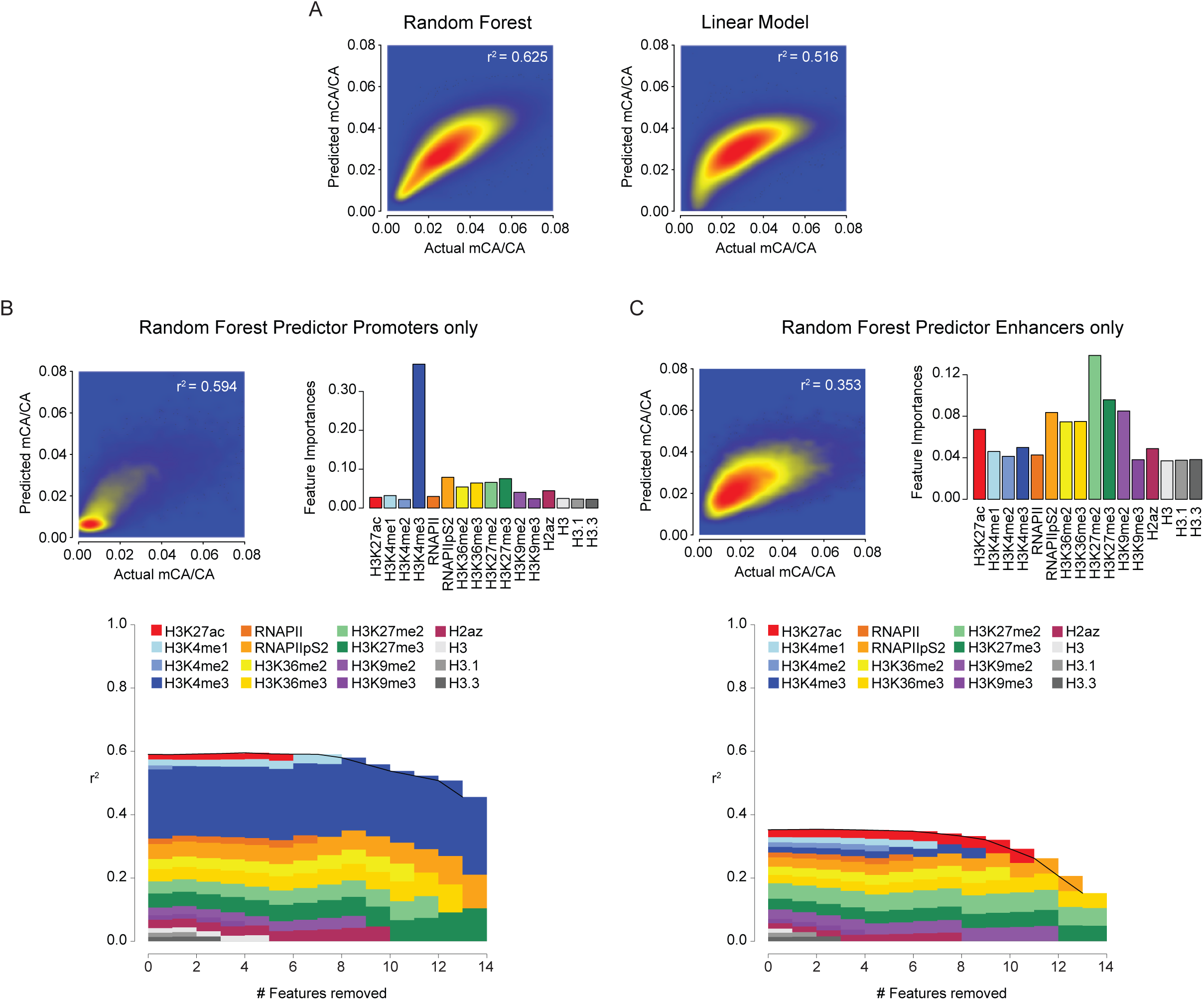
Analysis of mCA prediction by chromatin signals. Related to Figure 4. (A) Smoothscatter plot illustrating predictive accuracy of ChIP signals for mCA using a Random Forest algorithm (left) versus a linear model (right). Increased predictive power of the random forest suggests some non-linear or context-specific relationships between chromatin signals and mCA deposition (e.g. H3.3 intragenic and intergenic signals, Figure 4A). (B) Left, random forest classifier predicted mCA versus actual mCA levels at promoters (+/- 500bp from TSS center, see *methods*). Right, feature importance level of each chromatin signature to the predictive accuracy of mCA levels at promoters by the random forest classifier. Bottom, feature elimination analysis of chromatin signatures in the random forest classifier. (C) Left, random forest classifier predicted mCA versus actual mCA levels at enhancers (+/- 500bp from enhancer center, see *methods*). Right, feature importance level of each chromatin signature to the predictive accuracy of mCA levels at enhancers by the random forest classifier. Bottom, feature elimination analysis of chromatin signatures in the random forest classifier. Data are from wildtype cerebral cortex at 2- and 8-weeks. 2-week, n = 2-3 bioreplicates for H3K36me2, H3K4me1, H3K4me2, and DNMT3A ChIP-seq. 2-week RNA-seq data obtained from Lister et al., 2013. 2-week chromatin mark ChIP-seq and 8-week DNA methylation data obtained from Stroud et al., 2017.

**Figure S5.**
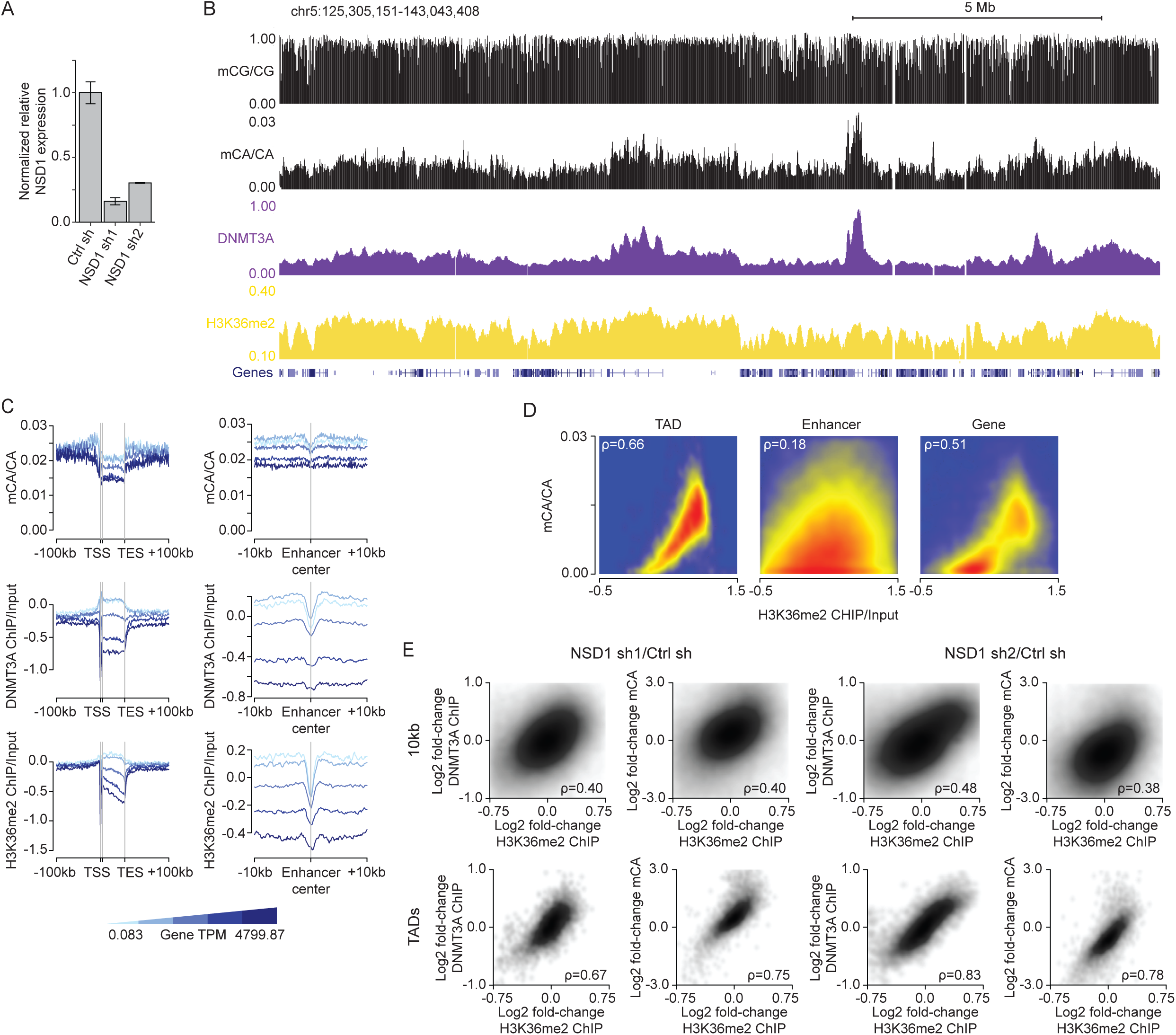
H3K36me2, DNMT3A, and mCA are correlated and concordantly disrupted upon NSD1 loss in primary cortical neurons (PCNs). Related to Figure 5. (A) qRT-PCR of Nsd1 over Actb normalized to shCtrl sample from two independent shNSD1 PCN samples (n = 2 per condition). (B) Genome browser view of DIV 12 ChIP-seq and DIV 18 DNA methylation from shCtrl PCNs. There is strong concordance of fluctuations in H3K36me2, DNMT3A, and mCA across the neuronal genome. (C) Aggregate mCA, DNMT3A/Input, H3K36me2/Input levels at genes and enhancers across quintiles of PCN gene expression. (D) Comparison of H3K36me2 ChIP/Input and mCA/CA at TADs, enhancers, and genes. (E) Comparison of changes in H3K36me2 and changes in DNMT3A (left) or changes in mCA (right) for each 10kb region (top) and TAD (bottom) across two independent shNSD1 samples. Data are from primary cultured neurons infected with shCtrl or shNSD1 after 1 day *in vitro* (DIV) and collected at DIV 12 and DIV 18 for ChIP-seq and WGBS, respectively. Per time point: n = 2-4 bioreplicates for H3K36me2, DNMT3A ChIP-seq, and DNA methylation. TADs were derived from Hi-C analysis of cortical neurons (Bonev et al., 2017).

**Figure S6.**
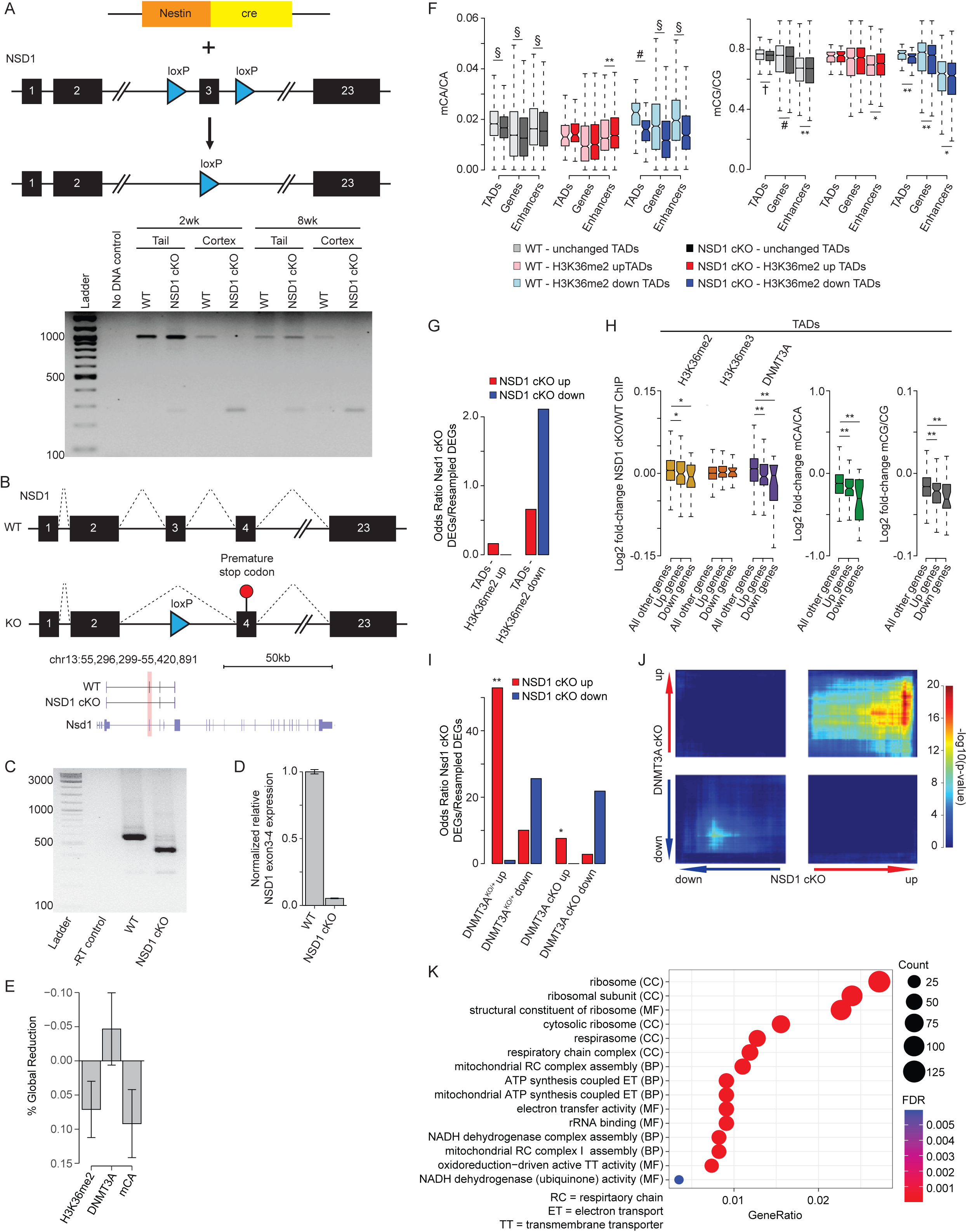
Conditional disruption of NSD1 in the brain leads to changes in H3K36me2, DNMT3A, and mCA that result in gene expression effects overlapping the DNMT3A Baf53-cKO mutant. Related to Figure 6. (A) Top, schematic outlining generation of the NSD1 cKO mutant allele. Bottom, genotyping PCR of the wildtype, flx/flx, and mutant alleles from tail or whole cortex at 2- and 8-weeks. (B) Top, schematic outlining mRNA splicing over excised exon 3 to generate shortened mRNA with premature stop codon and truncated NSD1 protein in the NSD1 cKO. Bottom, genome browser view of Nsd1 gene with sanger sequencing PCR products from NSD1 cKO and wildtype whole cortex mRNA illustrating successful excision of targeted exon. (C) PCR of NSD1 cKO and wildtype whole cortex cDNA illustrating successful excision of targeted exon from the major NSD1 isoform. (D) qRT-PCR of Nsd1 over Actb normalized to wildtype from NSD1 cKO and wildtype whole cortex mRNA (Per condition, n = 6). (E) Mean percent global reduction in H3K36me2, DNMT3A, and mCA observed in the NSD1 cKO using ChIP-Rx normalization (Orlando et al., 2014). (F) DNA methylation levels at TADs with significantly changed H3K36me2 (edgeR, FDR <0.1) and kilobase-scale genomic elements that reside within these TADs in NSD1 cKO and wildtype cerebral cortex. (G) Odds ratio of the overlap of NSD1 cKO significantly dysregulated genes with TADs with significantly changed H3K36me2. (Fisher’s exact test, observed versus background estimated by resampling). (H) Fold-changes of H3K36me2, H3K36me3, DNMT3A, and DNA methylation at TADs that overlap significantly dysregulated genes in the NSD1 cKO. (I) Odds ratio of the overlap of NSD1 cKO significantly dysregulated genes with DNMT3A^KO/+^ and DNMT3A Baf53-cKO significantly dysregulated genes (Christian et al., 2020; Clemens et al., 2019). (Fisher’s exact test, observed versus background estimated by resampling). (J) RRHO (Cahill et al., 2018; Plaisier et al., 2010) of all genes in the NSD1 cKO versus the DNMT3A Baf53-cKO (Clemens et al., 2019). (K) Top five Gene Ontology terms from “Biological Process”, “Molecular Function”, and “Cellular Components” from RRHO-determined “dd” (downregulated-downregulated) genes from NSD1 cKO and DNMT3A^KO/+^ using clusterProfiler (Wu et al., 2021; Yu et al., 2012). Data are from NSD1 cKO and control mouse whole cortex at two or eight weeks. Per time point: 2-weeks, n = 5 bioreplicates for H3K36me2, H3K36me3, DNMT3A ChIP-Rx. 8-weeks, n = 3 bioreplicates for DNA methylation, n = 6 bioreplicates for total RNA-seq. TADs were derived from Hi-C analysis of cerebral cortex (Clemens et al., 2019; Dixon et al., 2012). *p < 0.05, **p < 0.01, #p < 10^-5^, †p < 10^-10^, §p < 10^-15^.

